# Impact of structural sampling, coupling scheme and state of interest on the energy transfer in CP29

**DOI:** 10.1101/2023.01.25.525376

**Authors:** S. Petry, J. C. Tremblay, J. P. Götze

**Affiliations:** Department of Biology, Chemistry, Pharmacy, Freie Universität Berlin, Berlin, Germany; Laboratoire de Physique et Chimie Théoriques, CNRS-Université de Lorraine, 57070 Metz, France

## Abstract

The Q_y_ and B_x_ excitation energy transfer (EET) in the minor light harvesting complex CP29 (LHCII B4.1) antenna complex of *Pisum sativum* was characterized using a computational approach. We applied Förster theory (FRET) and the transition density cube (TDC) method estimating the Coulombic coupling, based on a combination of classical molecular dynamics and QM/MM calculations.

Employing TDC instead of FRET mostly affects the EET between chlorophylls (Chls) and carotenoids (Crts), as expected due to the Crts being spatially more challenging for FRET. Only between Chls, effects are found to be small (about only 0.1 EET efficiency change when introducing TDC instead of FRET). Effects of structural sampling were found to be small, illustrated by a small average standard deviation for the Q_y_ state coupling elements (FRET/TDC: 0.97/0.94 cm^−1^). Due to the higher flexibility of the B_x_ state, the corresponding deviations are larger (FRET/TDC between Chl-Chl pairs: 17.58/22.67 cm^−1^, between Crt-Chl pairs: 62.58/31.63 cm^−1^). In summary, it was found for the Q band that the coupling between Chls varies only slightly depending on FRET or TDC, resulting in a minute effect on EET acceptor preference. In contrast, the coupling in the B band spectral region is found to be more affected. Here, the S_2_ (1B_u_) states of the spatially challenging Crts may act as acceptors in addition to the Chl B states. Depending on FRET or TDC, several Chls show different Chl-to-Crt couplings. Interestingly, the EET between Chls or Crts in the B band is found to often outcompete the corresponding decay processes. The individual efficiencies for B band EET to Crts vary however strongly with the chosen coupling scheme (e.g., up to 0.29/0.99 FRET/TDC efficiency for the Chl *a*604/neoxanthin pair). Thus, the choice of coupling scheme must involve a consideration of the state of interest.

## Introduction

The photosynthetic machinery of plants is a highly complex system of several modules, namely protein-pigment complexes, acting, sometimes simultaneously, as antennae, conductors and reaction centers (RCs).[1-5] Quantitatively, most modules are light-harvesting complexes (LHCs), which acquire the bulk of the photonic energy. Although detailed knowledge regarding molecular photosynthetic processes is established, as they have been studied since mid-last century, they are still not fully understood.^4–12^ Especially the energy pathways are difficult to identify as there are plenty of possibilities. Due to their evolutionary origin, they do not need to follow a straightforward logic akin to chemical engineering, which might make their mechanistic principles very unintuitive. Still, several possible pathways have been investigated from both the experimental and theoretical perspective.^13–21^.

CP29 is a minor light-harvesting antenna complex^22,23^ of the photosystem II (PSII) supercomplex, which is present in higher plants and green algae. CP29 is located between the core complex and a moderately bound Light Harvesting Complex II (LHCII) with proximity to a strongly bound LHCII and the minor light-harvesting antenna complex CP24 (see Figure 1). All modules of the PSII supercomplex contain two types of chromophores, chlorophylls (Chls) and carotenoids (Crts). CP29 contains ten Chl *a*, four Chl *b* as well as 3 Crts, namely neoxanthin, lutein and violaxanthin.^24,25^ The Chl *a* are the photosynthetically active pigments in the RCs, thus determining many biophysical properties of Chl-containing complexes.^26–31^ Their role is harvesting sunlight, transferring energy and, in the RC, charge separation.^32–35^ Chls (*i*.*e*., not only Chl *a*) have two absorption bands in the UV/Vis region, namely the Q band (comprising the electronic Q_y_ and Q_x_ states) and the B (“Soret”) band, which contains at least two electronic states as well (B_x_ and B_y_).^36^ As the Q band is fuels the biological function of charge separation, many investigations have been focused on the Q band.^37–41^ The role of the Crts has been attributed mainly to photoprotection and supporting the light absorption of the Chls; the latter also being the supposedly primary role of Chl *b*.^20,42,43^ The post-absorption processes in the Chls and Crts for the blue spectral region are currently mostly explained by fast internal conversion (IC) or vibrational relaxation (VR).^44^ There are however reports of energy transfer between Crts to Chls involving the ultraviolet/blue spectral region (350-550 nm).^17,45–49^

**Figure 1:**
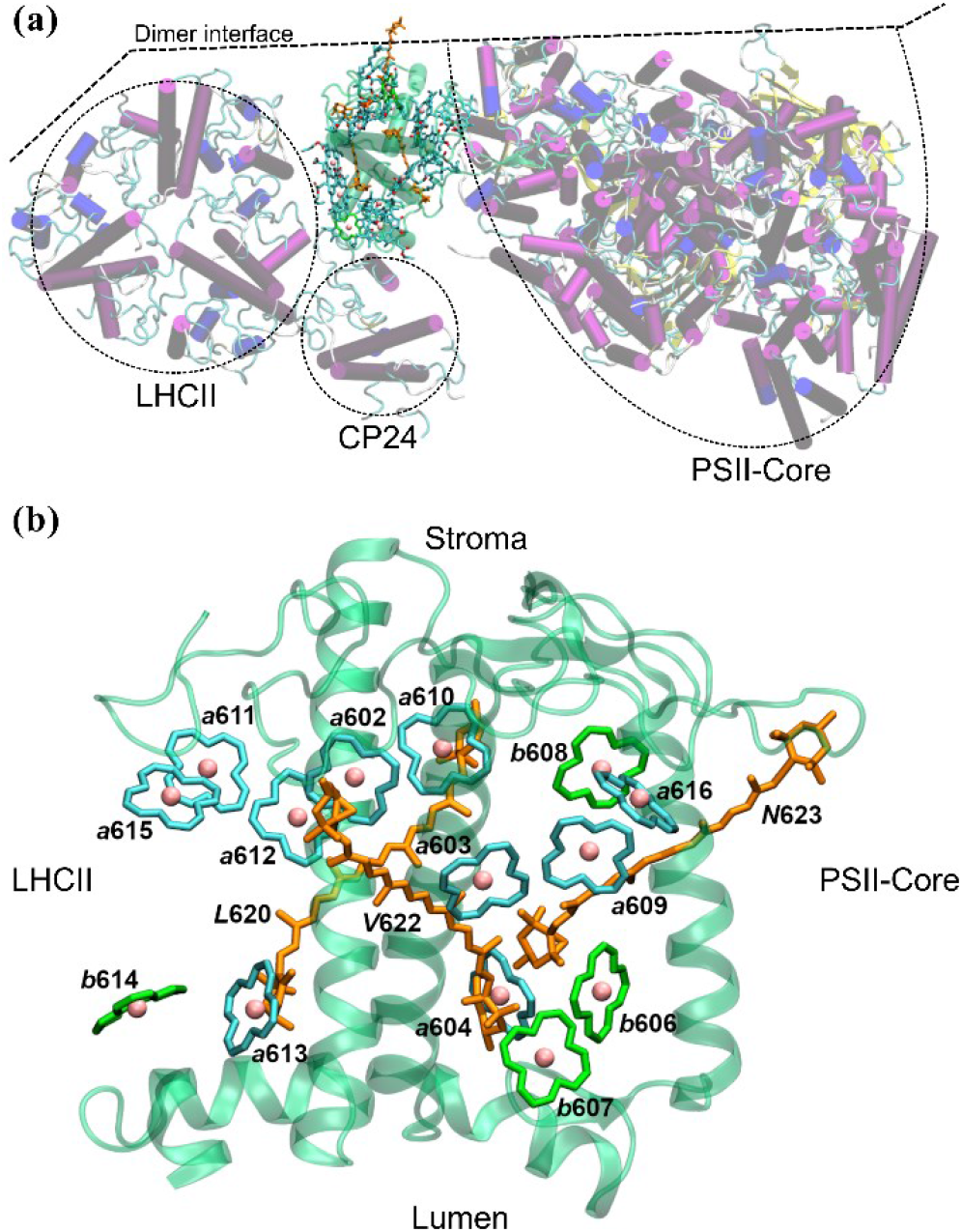
(a) Arrangement of CP29 within a photosystem II monomer viewed from the stromal side of the membrane, encircled are the moderately bound LHCII (M-LHCII), the antenna complex CP24 and the PSII core complex (PSII-Core). (b) CP29 structure with the protein backbone (turquoise) and the chromophores Chl a/b (cyan/green) core ring atoms and the Crts (orange) violaxanthin (V622), neoxanthin (N623) and lutein (L620). The chromophores are labelled for identification.

CP29, as well as other LHCs such as LHCII, have been attributed photoprotective functions (non-photochemical quenching, NPQ). The molecular origin of NPQ is not fully understood, as many simultaneous events may result in similar observations. There are plenty of postulated mechanisms, especially for the LHCII.^50–54^ Many, but not all of these proposals require Crts.

In protein-pigment complexes like CP29, the protein part organizes the position of the chromophores and modifies the optical properties by defining the electrostatic environment.^55^ Unsurprisingly, the direct Mg-ion ligand was found to be potentially determining for the Chl absorption. ^56^ In this earlier study, we found that for CP29, molecular dynamics (MD, for sampling) followed by a coupled quantum mechanics/molecular mechanics (QM/MM, for optimization and time-dependent density functional theory calculation (TD-DFT)) is necessary to obtain a proper ordering of Chl *a*/*b* states. The study focused on the environmental protein matrix effect for the coupling between Chls using a basic Förster resonance energy transfer (FRET) model. The approach had to be restricted to the Chls, neglecting the Crts, because the spatially extended Crt transition densities exaggerated the error from the point dipole approximation (PDA). *I*.*e*., the transition dipole moments (TDMs) within PDA could no longer be assigned to a certain spatial point for Crts, making it impossible to define the intermolecular distances. Furthermore, the calculation of rates in FRET theory is strongly depending on the quality of the computed spectra (see FRET details in the next section). The performance of quantum chemical methods depends however on the chromophore type, and Chls are much easier to tackle than Crts.^57^

As an alternative for FRET, and thus the PDA, the transition density cube (TDC) method estimates the *full* Coulomb coupling. This is in contrast to FRET, which only covers dipole-dipole coupling. TDC may possibly also provide the total electronic coupling as well, by taking the transition density overlap into account as shown by Scholes et al. ^58^. Because the actual spatial shape of the transition densities is used, it should be valid for any molecular shape, making the PDA unnecessary.

In this article, we address the issues of (i) balanced, non-PDA representation for Chl-Chl and Crt-Chl coupling and (ii) potential involvement of B band states in Crt-Chl interactions. We start by exploring the Coulomb coupling of CP29 chromophores, comparing FRET and TDC qualitatively and quantitatively, where appropriate. We include the Q_y_ as well as the B_x_ state interactions and the respective differences. Both Q_y_ and B_x_ are the lowest energy states of their respective optical bands, and thus the ones most likely to act as energy donors. The Coulomb coupling of the Crts with the B_x_ state of Chls is evaluated, as those will be shown below to provide significant coupling regardless of the employed coupling scheme.

## Methodology

### Computational Details

#### Geometries and QM/MM model

From a 100 ns MD simulation performed in a NPT ensemble at T = 300 K using leap-frog integration, 11 snapshots were taken, with an interval of 5 ns to obtain the structurally uncorrelated initial structures for the QM/MM ensemble (for more details see our previous study ^56^). The initial structures were reoptimized with QM/MM to perform TD-DFT in QM/MM setup to obtain the spectral properties. Consequently, Chl calculations and properties were also identical to the previous study. Note that due to our model only including CP29, Chl *a*616, which is at the interface to the PSII CP47 antenna, was not considered. The QM/MM calculations were performed with the gmx2qmmm interface^59^ using Gaussian 16^60^ and the GROMACS simulation package 2018.6^61^. All QM and QM/MM calculations employed the CAM-B3LYP functional with the 6-31G* basis set; the force field was in all cases AMBER99SB*-ILDNP.

Compared to our earlier work, Crts were added here to the chromophore ensemble. The corresponding QM-layers consisted solely of the Crt in question, as no strong binding partners were found at any point during the MD for any of the Crts. The inner (interacting) layer included all atoms within a range of 4 nm with respect to the nearest atom of the Crt. The active region consisted of all protein residues, lipids, solvent molecules and ions containing atoms in a rough van-der-Waals distance (0.3 nm) of the QM layer. The active region was optimized using a steepest descent algorithm (force tolerance: 100.0 kJ/mol/nm, step size 0.01 nm, 3 nm radius of non-bonded partner list). To further reduce computational costs, all solvent molecules with a distance larger than 2 nm to an atom of the QM layer were automatically removed from the system beforehand

#### State energies

After QM/MM optimization, the structures were used for TD-DFT^62^ calculations of the first twelve lowest states to simulate vertical absorption spectra. The Chl Q_y_ state was always taken as the lowest Chl excited state. The B_x_ state as the lowest state of the B band with an oscillator strength larger than 0.1. Other states were only computed for diagnostic purposes.

In the present study, excited state energies are thus represented via the vertical excitations (*E*_vert_). *E*_vert_ was used regardless of excitation (absorption) or de-excitation (emission). This neglects the reorganization energy from excited state relaxation following absorption. While this approximation is fairly good for Chls displaying a small reorganization energy of only 0.2-0.3 eV, and virtually no experimental Stokes shift^63,64^, it is not suitable for Crts with a reorganization energy of 0.4-0.5 eV (Figure 2)^57^, sometimes higher.^65^ To account for this, only Chl emissions are considered in this article; *i*.*e*., Crts only act as acceptors in our model due to the uncertainty of the emission wavelength compared to Chls.

**Figure 2:**
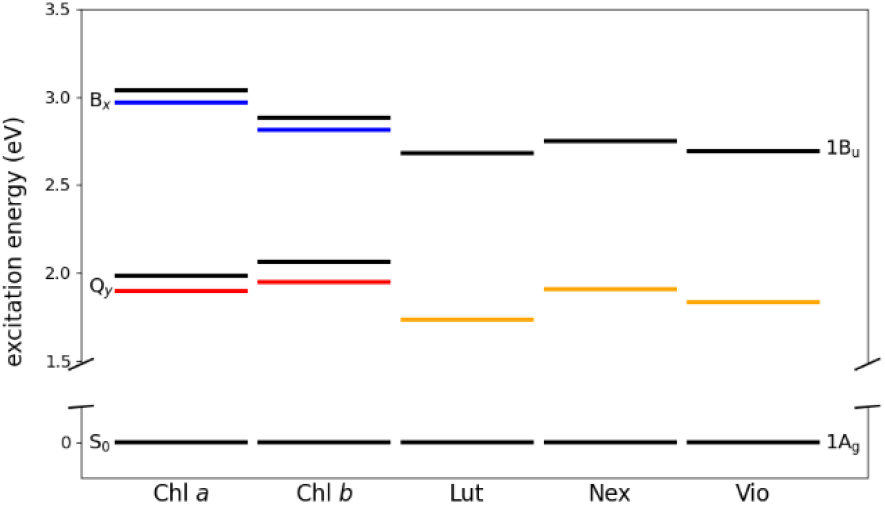
Computed Stokes shifts of investigated chromophores, data from ^57^. Jablonski diagrams displaying E_vert_ for the indicated states (Q_y_, B_x_ and 1B_u_) in black (after TD-DFT nuclear relaxation of the ground state), for Chl a, Chl b, lutein (Lut), neoxanthin (Nex) and violaxanthin (Vio). The colored lines indicate E_vert_ for the respective state after excited state optimization using DFT/MRCI: Q_y_ state (red), B_x_ state (blue) and the 1B_u_ state (orange). The difference between black and colored lines is the computed, vertical, Stokes shift. Other states of Chls and Crts (such as Q_x_ for Chls or the dark 2A_g_ state of Crts) omitted for clarity.

### Coupling Analysis

#### Förster model

In FRET, the coupling of two separated chromophores is described using only inter-chromophore distance as well as the TDMs or derived quantities.^66–70^

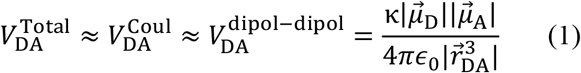

Here, 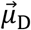 and 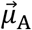 are the TDMs of the donor (D) and acceptor (A), 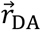 is the vector connecting the two point sized TDMs and *ϵ*_0_ is the vacuum permittivity. The orientation factor κ defined as:

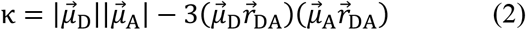

This requires the definition of a molecular center for the two molecules. Consequently, the PDA produces errors for molecules with extended or asymmetric transition densities, introducing a larger error for Crts.^71^ Further, at small 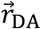, orbital overlap may occur which is also not accounted for FRET.

#### TDC

The TDC method, developed by Krueger et al.^66^, estimates the Coulomb coupling by using the transition densities of the donor and acceptor excited states. Computationally, the transition densities are represented in TDC as three-dimensional cubes representing a set of finite-sized volume elements *V*.

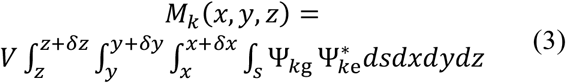

With Ψ_*kg*_ being the wavefunction of an initial state and Ψ_*ke*_ the wavefunction of a final state of the chromophore *k*. The Coulomb coupling 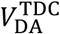 of the donor and acceptor states in TDC is then approximated as the sum over the Coulomb interactions of these spatial elements.

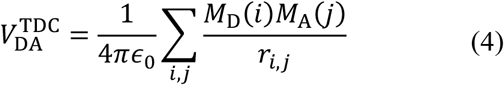

For the calculation of the transition density cubes, the ORBKIT package was used.^72–74^ The accuracy of the method is limited by the accuracy of the wavefunction and the size of the volume elements (grid size). Further, within TDC we are still restricted to the Coulomb coupling, not taking orbital overlap effects into account. The other downsides of FRET are however not present in a spatially inclusive method such as TDC. Unfortunately, the resulting transition densities require scaling to yield observables that can be compared to experiment. To do so, the transition densities are scaled such that the calculated transition multipoles match the magnitude of the TD-DFT calculated TDMs.^66^ This approach is common in TDC and also related methods, such as TrESP (transition charges from electrostatic potential).^75–77^

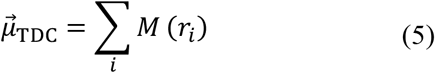

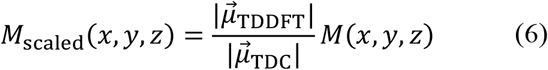

These scaled elements can then be used in Eq. (4) to calculate Coulomb couplings.

#### Rates

Coupling elements from FRET or TDC were used to estimate the EET (excitation energy transfer) rates *Γ*_DA_ of two interacting chromophores using Fermi’s golden rule:

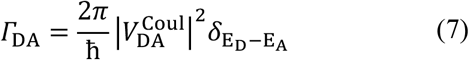

Here, the delta function (which is similar, but not identical to the spectral overlap *J* in the FRET approach used for experiments)^78^ is approximated with the integral of the product normalized fluorescence spectrum *F*_D_(*v*) of the donor and the normalized absorption spectra *A*_A_(*v*) of the acceptor.^79^

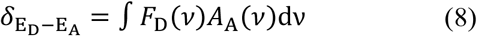

Where:

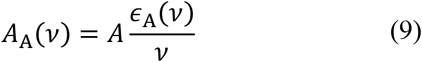

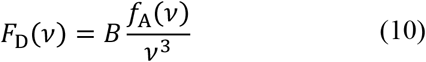

with A and B as normalization factors, *ϵ*_A_ as an absorption spectrum and *f*_A_ as an emission spectrum. This results in 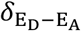 requiring a function describing the absorption and emission spectra. This is provided by Lorentzian functions *L*(*E*) centered at the site energy of the donor *E*_D_ and the acceptor *E*_A_. The choice of a full width at half maximum *γ* = 2000 cm^−1^ corresponds to the width of an experimental Q_y_ state at room temperature). ^63^

Neglecting the reorganization energy from excited state relaxation as noted above results in *ϵ*_A_(*v*) = *f*_A_(*v*). However, the normalized fluorescence spectrum (Eq. (10)) differs from the normalized absorption spectra (Eq. (9)) due to the factors 1/*v* and 1/*v*^3^. Further, a direction of EET will still appear between two identical chromophores because of the different CP29 site energies. In any case, for the purpose of this article, comparing FRET and TDC, the exact state energies are not relevant if 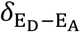 is the same for both approaches.

#### Förster radii for TDC and EET efficiencies

In FRET theory, the Förster radius or FRET radius *R*_0_ is defined as the distance at which the rates of EET and excited state decay are identical (*Γ*_DA_(*r*_DA_ = *R*_0_) = 1/*τ*_f_). Lifetimes were taken from experiment, see Table 1.

Conversely, FRET radii can be calculated using the rates of any EET approach, also TDC.

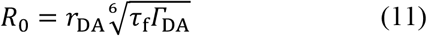

The EET efficiency *E*_EET_ of a transfer is defined as the ratio of the EET rate *Γ*_DA_ and the fluorescence rates *Γ*_f_, the sum of all energy transfer rates ∑*Γ*_DA_ and other non-radiative decays ∑ *Γ*_nr_. The experimental donor fluorescence lifetime in absence of an acceptor *τ*_f_ is the sum of all non-FRET decay lifetimes, leading to Eq. (12).

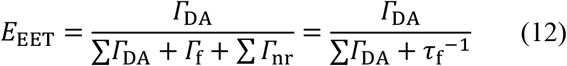

In comparison to that, the pairwise EET efficiency 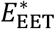 is the ratio of a single EET rate *Γ*_*DA*_ and the inverse experimental lifetime *τ*_f_^−1^ (Eq. (13)).

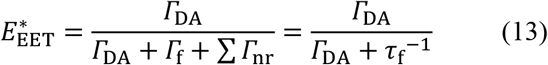

**Table 1:** Experimental lifetimes of the Q_y_ and B_x_ states for Chl a, b and the lifetime of the S_2_ state of Crts. ^80–82a^ Average of a reported 100-250 fs range.

| Chromophore | $Q_y$<br>$\tau_f/\text{ns}$ | $B_x$ (Chl) or $S_2$ (Crt)<br>$\tau_f/\text{fs}$ |
| --- | --- | --- |
| Chl *a* | 6.3 | 175<sup>a</sup> |
| Chl *b* | 3.2 | 58 |
| Crt<sub>s</sub> | - | 163 |

## Results and discussion

We split the results in two parts, separately discussing the Q band and the B band. The Q band EET is represented here by Q_y_ state interactions, and the B band EET conversely by B_x_ state interactions. The first part, which is focus of several studies^55,83,84^ shows the reliability of our calculations. The B band part is a novel analysis, as the B band has rarely been considered as a viable EET band in the context of Chls.^34,85–87^

### Q band

We start with an analysis of Chl-Chl pairs, Chl *a*/*a, a*/*b* and *b*/*b* for the first excited state (the Q_y_ state). The full information of average rates and efficiencies can be found in the supporting information (SI). Due to the large number of possible Chl pairs, we discuss only sets of pairs with increasing spatial distance.

At this point it should be re-emphasised that *V*_FRET_ and *V*_TDC_ are in both directions identical due to the neglect of reorganization energy. Only the rates and the rate-dependent values are different for the reverse process due to a slightly different 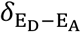 (exchanging *F*_D_ and *A*_A_ in Eq. (8)). Therefore, the EETs between Chls *b* (donor) and Chls *a* (acceptor) are not discussed in detail as *V*_FRET_ and *V*_TDC_ are given from the *a*→*b* case. The rates are similar, but slightly lower, due to the blue shift of Q_y_ for Chl *b*.

#### EET for Q_y_ between Chls a--a

Table 2 (row 1-4) shows *V*_FRET_ and *V*_TDC_ of Chl *a*612 (donor) and four acceptor Chls *a* with increasing distance, namely *a*611, *a*613, *a*603, *a*609 with an average *r*_DA_ of 9.2, 18.0, 22.2, 29.0 Å, respectively. The choice for *a*612 as example was made due to the comparable distances with respect to the other *a*-*b* or *b*-*b* cases of the other sections; *a*612-*a*611 also shows the strongest coupling. Table 2 also shows the calculated *R*_0_ resulting from *V*_FRET_ and *V*_TDC_ as well as κ^2^. In the SI, Figure S1 shows the dynamical evolution of the *V*_FRET/TDC_ and the corresponding *R*_0_.

**Table 2:** Comparison of the calculated average distances (*r*_DA_), κ^2^ values, Coulomb couplings of the FRET (*V*^FRET^) and TDC-scheme (*V*^TDC^) and the corresponding average FRET radii for the Q_y_ state.

| D-A pair | $r_{DA}/\text{\AA}$ | $\kappa^2$ | $V^{FRET}/\text{cm}^{-1}$ | $V^{TDC}/\text{cm}^{-1}$ | $R_0^{FRET}/\text{\AA}$ | $R_0^{TDC}/\text{\AA}$ |
| --- | --- | --- | --- | --- | --- | --- |
| *a612-a611* | 9.2±0.1 | 2.05±0.12 | 225.55±30.03 | 163.98±26.43 | 57.15±1.36 | 51.41±3.22 |
| *a612-a613* | 18.0±3.5 | 0.04±0.04 | 3.39±2.46 | 2.89±2.65 | 26.08±5.86 | 22.77±7.22 |
| *a612-a603* | 22.2±0.1 | 0.06±0.07 | 2.30±1.26 | 4.68±1.24 | 29.11±5.07 | 37.68±3.33 |
| *a612-a609* | 29.0±0.5 | 1.30±0.23 | 5.47±0.51 | 4.38±0.70 | 52.23±2.20 | 48.38±2.55 |
| *a604-b606* | 8.1±0.1 | 0.70±0.11 | 129.25±27.62 | 76.57±20.74 | 40.94±2.82 | 34.27±3.25 |
| *a604-b607* | 14.0±0.2 | 0.02±0.03 | 3.89±2.80 | 7.69±5.03 | 20.76±4.91 | 26.64±6.02 |
| *a604-b608* | 17.7±0.0 | 0.24±0.07 | 7.37±1.28 | 0.76±0.68 | 34.62±1.77 | 14.62±5.25 |
| *a604-b614* | 29.2±1.0 | 0.44±0.25 | 2.03±0.97 | 2.08±0.74 | 36.24±6.34 | 37.16±4.73 |
| *b606-b607* | 9.8±0.4 | 0.07±0.07 | 14.53±8.51 | 18.78±14.56 | 20.91±3.55 | 21.74±7.36 |
| *b606-b608* | 16.6±0.1 | 0.22±0.07 | 5.46±1.87 | 3.42±2.43 | 26.51±2.47 | 21.04±6.42 |
| *b606-b614* | 36.3±0.9 | 1.03±0.24 | 1.17±0.29 | 0.92±0.36 | 34.52±3.17 | 31.44±4.88 |

Chls can obviously also vary in their strength of coordination to the local protein matrix. This has a direct effect on the variance in *r*_DA_. As an example, of the cases displayed, Chl *a*613 (Table 2, second row; also Figure S1B) is only transitionally bound to glutamine (7 out of 11 frames). It is thus structurally more dynamic than the other cases and *r*_DA_ becomes 18.0±3.5 Å. The other three pairs in Table 2 have a standard deviation of 0.52 or lower. The comparatively high standard deviation of *r*_DA_ between Chl *a*612 and Chl *a*613 indicates a weak bond and thus a high flexibility, which is in a good agreement with a previous theoretical study ^54^.

For each D-A pair, *V*_FRET_ and *V*_TDC_ in Table 2 is in the same order of magnitude (FRET/TDC, in cm^−1^): For *a*611, 226±30/164±26, for *a*613 3±3/3±3 for *a*603 2±1/5±1 and for *a*609 6±1/4±1. As TDC and FRET are not strictly independent due to the rescaling of the TDMs in TDC, this was to be expected. However, there is no straightforward statement possible with regards to *V*_TDC_ and *V*_FRET_ being always larger or smaller; if there is a correlation, it is more complex than that and not directly obvious from our data. This non-systematic relationship has already been observed before and is thus consistent with our expectations. ^88^

The main factor affecting *V*_FRET_ and *V*_TDC_ is the distance, which is not surprising (Eq. (1)). For example, the computed *V*_FRET_ and *V*_TDC_ of Chl *a*612 and *a*611 are strong with an average of 226±30/164±26 cm^−1^ (FRET/TDC), due to the small *r*_*DA*_ and a comparatively high average κ^2^ of 2.05. The weak couplings for the other Chl pairs are a due to low average κ^2^ values (0.04 and 0.06 for *a*612-*a*613 and *a*612-*a*603). Although, Chl *a*609 is with 29.0 Å the furthest Chl of the three “weak” Chls, the average *V*_DA_ is somewhat stronger with 5.47/4.38cm^−1^ due to an average κ^2^ of 1.30. The *R*_0_ values for FRET/TDC for the closest Chl pair (*a*612-*a*611) is around 57.2±1.4 / 51.4±3.2 Å and for the furthest (*a*612-*a*609) 52.2±2.2 / 48.4±2.6 Å, which is in a good agreement with the experiment.^89^ For the other displayed Chl *a-a* pairs, the 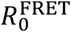 are 26.1±5.9/29.1±5.1 Å (*a*612-*a*613/*a*612-*a*603) and the 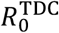 22.8±7.2 / 37.7±3.3 Å (*a*612-*a*613/*a*612-*a*603), resulting from the unfavorable κ^2^.

In terms of rates, for Chl *a*612 and *a*611, 9.4/5.2 ps^−1^ is obtained, yielding a near-unity pairwise EET efficiency (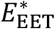 above 0.99) for both methods (see Table S3 and S5 in the SI). Conversely, for the other Chl *a-a* D-A pairs, rates are much lower, and hence, 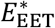 is reduced. However, the “weakest” *V*_DA_ of *a*612-*a*603 still refers to 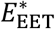 of 88% for FRET and 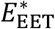 of 95% for TDC. This means that at Chl-Chl distances found within CP29, even a slightly non-zero κ^2^ will lead to a high, sometimes near-unity 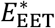 for the Q band.

Considering all possible Chl *a*--*a* pairs in CP29, not only those from Table 2 with Chl *a*612 as donor, the lowest calculated FRET/TDC rate is 0.10/0.11 ns^−1^ (*a*613-*a*609 pair, with an average *r*_*DA*_ of 32.7 Å). While the average distance of another pair (*a*615-*a*609) is higher, (34.7 Å), the corresponding κ^2^ is higher with 0.59 (*a*615-*a*609) compared to 0.04 (*a*613- *a*609).

However, even the weakest rates of the *a*613-*a*609 pair still correspond to 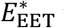 values of 33/27 % (see Figure 5). This is the only coupling with an efficiency below 50% for any Chl *a*--*a* pair using the FRET scheme. For TDC, only *a*611-*a*603 (48%) and *a*603-*a*611(49%) show 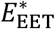 below 50% as well. This means that all Q band EET rates are competitive compared with the Q band decay (IC and fluorescence; 1/*τ*_*D*_). However, the CP29-wide *E*_EET_ values show acceptor preferences for each donor (see Table S4 and S6 in the SI, each donor has only a few cases for network *E*_*EET*_ > 0). While this of course might indicate a steering of EET, likely to the spatially closest acceptor, it also shows that the Q band EET network is very robust: The Chls are so close in CP29 that the loss of individual Chls (failures in complex assembly, mutations or damage etc.) would not affect overall Q band EET efficiency, only the preferred acceptor

#### EET for Q_y_ between Chls a--b

Table 2 (rows 5 to 8) shows *V*_DA_ of Chl *a*604 (selected example donor) and all four CP29 Chls *b* as acceptors, namely *b*606, *b*607, *b*608, *b*614, with an average *r*_*DA*_ of 8.1, 14.0, 17.7, 29.2 Å, respectively. For the investigated Chl pairs, *V*_DA_ of *a*-*b* is generally less than *V*_DA_ of *a*-*a*, which is correlated with the weaker Q band oscillator strength of Chl *b* compared to Chl *a*.

The closest Chl *a*-*b* pair *a*604-*b*606 shows a strong average coupling of 129.3±27.6/76.6±20.7 cm^−1^ (FRET/TDC), with the average 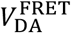 being 40% higher; here, FRET apparently overestimates the coupling drastically. The average κ^2^ of all shown *V*_DA_ are relatively low with 0.70, 0.02, 0.24 and 0.44 (*a*604- *b*606, -*b*607, -*b*608 and -*b*614) considering that κ^2^ is 2/3 for freely moving chromophores.^90^ The FRET/TDC rate is 2.9/1.0 ps^−1^ for the closest *a*-*b* pair *a*604-*b*606 and the rates are 4.0/13.8, 9.5/0.2, 0.9/0.8 ns^−1^ for -*b*607, -*b*608, -*b*614 respectively (note the change in order of magnitude; these rates are much slower than the other presented rates in the article). Even for the weakest *V*_DA_ *a*-*b* pair, the rates of 0.9/0.8 ns^−1^ correspond to 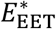 values of 71/77%. Hence, the 40% overestimation of 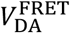 for the strongest *a*-*b* (*a*604-*b*606) pair noted above is not relevant for 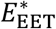, as both coupling values will result in near-unity values. Still, for the full set of competing EET processes (see Table S1 and S2 in the SI), this also makes only a minor difference of shifting from 0.94 *E*_*EET*_ in FRET down to 0.86 *E*_*EET*_ in TDC. So far, differences between FRET and TDC do not qualitatively, and only minutely quantitatively, change the EET picture.

Note that the *V*_DA_ values were not explicitly discussed in this section due to the limited insight compared to the insights from discussing the EET efficiencies. For one case, however, we need to highlight the couplings again as there is a much more drastic deviation for FRET vs. TDC. Chl *b*606 shows a considerable discrepancy comparing 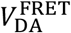 and 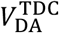 with 7.4 and 0.8 cm^−1^, respectively. However, this is only the case for the Q_y_ states and not for the B_x_, as discussed below. The Chl *b*606 and *b*608 are close to glutamate groups, which are connected to the Mg ions via hydrogen bonds through one or two bridging water molecules. The observed discrepancy is an effect of the structural sampling because the water molecule coordinating to the Chl *b*608 forms a hydroxide in four frames. Due to the electronegativity, the transition densities/TDMs of the Q_y_ state of Chl *b*608 become thus more delocalized (data not shown). FRET thus likely overestimates the coupling due to the PDA being less applicable here.

#### EET for Q_y_ between Chls b--b

The Chls *b*606, *b*607 and *b*608 are on the CP29 side that is facing the PSII core, while Chl *b*614 is f0acing the LHCII. With *r*_DA_ = 9.8 Å, Chl *b*606/*b*607 is the closest Chl *b* pair, located on the lumenal side of CP29. The spatially closer Chls *b-b* pairs are unfavorably orientated for coupling, which is represented by the low κ^2^ values (see Table 2, lower part) of 0.07(*b*606/*b*607), 0.22(*b*606/*b*608); the only pair with a favourable κ^2^ is *b*606-*b*614, with a distance of 36.3 Å and thus still the weakest *V*_DA_. However, for that weakest *b-b* pair the rate is 0.3e^−3^/0.2 e^−3^ ps^−1^ and 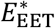 is 43/33% (FRET/TDC). This underlines the above argument that EET is efficient in the Q band, even at significant distances and unfavourable orientation.

#### Q_y_ network

As mentioned above, we restrict our discussion to select examples. The full FRET and TDC Q band network *E*_EET_ can be found in the SI. We find that for every Chl chromophore, a preferred Chl acceptor exists, which is mostly due to their proximity. Slight variations arise due to low κ^2^ for some spatially close pairs: For example, the distance of Chl *a*602 and *a*603 is 12.2±0.1 Å but as κ^2^ is 0.13±0.08, the corresponding *E*_EET_ is only 0.06/0.02 (donor: *a*602) or 0.07/0.03 (donor: *a*603) for FRE T/TDC, respectively. Chl *a*602 instead donates much better to *a*615, with *E*_EET_ of 0.82/0.94, despite that pair being 12.9±0.1 Å apart. Nevertheless, for the full CP29 Q band network, IC shows is below 0.01 for every Chl, implying that IC is not competing significantly with EET on the ps^−1^ timescales. Figure 4 from the Q/B comparison below shows that the FRET and TDC method show qualitative the same transfer rates. The only outlier is pair *a*604*-b*608 with a factor of about 50, which is discussed above as a result of PDA being less applicable for Chl *b*608.

### B band

In this section, we analyze the coupling of the B_x_ states for the same Chls as above. Additionally, the S_2_ states of the three Crts of CP29 will be considered as potential acceptors. While we showed above that EET between Q_y_ states has no significant competing process (*i*.*e*., IC is much slower than EET), we will see that this can also be the case for the B_x_ state. Further, for the B band, we mostly omit the discussion of 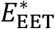 (see Figure 5) and focus on the network *E*_*EET*_ for the sake of brevity (see Table S9, S10, S11 and S12 in the SI). In the SI, Figure S2 shows the dynamical evolution of the *V*_FRET/TDC_ and the corresponding *R*_0_.

Due to the higher TDMs of the B states compared to the Q states, the couplings of the B_x_ states are generally stronger, for both FRET and TDC. The corresponding rates are thus consequently stronger as well (see Table S7 and S8 in the SI), but IC for the B band is also much faster than for Q (see Table 1). It will be seen below in which cases the IC processes may be outcompeted by B band 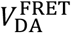 or 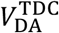.

#### EET for B_x_ between Chls a--a

The pairs *a*612-*a*611, *a*612-*a*613 and *a*612-*a*603 show 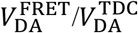 of 305.3/362.2, 36.4/35.6 and 11.5/7.8 cm^−1^ (see Table 3), which is stronger compared to the Q_y_ state coupling. *V*_DA_ of *a*612-*a*609 is lower in the B band than in the Q band, with 1.5/1.9 cm^−1^ (B) vs. 5.5/4.4 cm^−1^ (Q). Expressed in rates, this yields 19.6/27.3, 0.7/2.25 ps^−1^ for *a*612-*a*611, *a*612-*a*613 and rates below 0.1 ps^−1^ for *a*612-*a*603 and *a*612-*a*609. This is mainly due to the favourable κ^2^ of the former two pairs. While 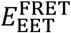 of the strongest *V*_DA_ Chl pair *a*612-*a*611 is with 18% in the same magnitude as the internal conversion with 16%, 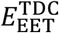 is with 42% higher compared with the IC with 27%.

**Table 3:** Comparison of the calculated average distances (*r*_DA_), κ^2^ values, Coulomb couplings of the FRET (*V*^FRET^) and TDC-scheme (*V*^TDC^) and the corresponding average FRET radii for the B_x_ state.

| D-A pair | $r_{DA}/\text{\AA}$ | $\kappa^2$ | $V^{\text{FRET}}/\text{cm}^{-1}$ | $V^{\text{TDC}}/\text{cm}^{-1}$ | $R_0^{\text{FRET}}/\text{\AA}$ | $R_0^{\text{TDC}}/\text{\AA}$ |
| --- | --- | --- | --- | --- | --- | --- |
| *a612-a611* | $9.2 \pm 0.1$ | $0.93 \pm 0.08$ | $305.29 \pm 68.55$ | $362.16 \pm 71.57$ | $12.22 \pm 3.59$ | $12.96 \pm 3.85$ |
| *a612-a613* | $18.0 \pm 3.5$ | $1.00 \pm 0.58$ | $36.43 \pm 19.70$ | $35.57 \pm 23.46$ | $10.03 \pm 2.01$ | $9.79 \pm 1.96$ |
| *a612-a603* | $22.2 \pm 0.1$ | $0.36 \pm 0.20$ | $11.48 \pm 5.24$ | $7.83 \pm 5.10$ | $9.56 \pm 2.33$ | $8.11 \pm 2.01$ |
| *a612-a609* | $29.0 \pm 0.5$ | $0.06 \pm 0.06$ | $1.49 \pm 1.25$ | $1.90 \pm 1.39$ | $5.28 \pm 2.23$ | $6.50 \pm 1.91$ |
| *a604-b606* | $8.1 \pm 0.1$ | $0.43 \pm 0.19$ | $355.24 \pm 80.10$ | $311.72 \pm 64.52$ | $9.72 \pm 0.86$ | $9.31 \pm 0.74$ |
| *a604-b607* | $14.0 \pm 0.2$ | $0.07 \pm 0.09$ | $22.09 \pm 15.09$ | $29.24 \pm 14.50$ | $7.17 \pm 4.03$ | $8.10 \pm 3.87$ |
| *a604-b608* | $17.7 \pm 0.0$ | $1.41 \pm 0.28$ | $57.10 \pm 16.62$ | $41.03 \pm 14.96$ | $11.94 \pm 1.75$ | $10.63 \pm 1.58$ |
| *a604-b614* | $29.2 \pm 1.0$ | $0.95 \pm 0.47$ | $10.78 \pm 3.26$ | $13.57 \pm 3.97$ | $11.33 \pm 3.23$ | $12.22 \pm 3.38$ |
| *b606-b607* | $9.8 \pm 0.4$ | $0.22 \pm 0.12$ | $165.48 \pm 74.56$ | $120.62 \pm 121.38$ | $7.58 \pm 1.68$ | $6.53 \pm 2.68$ |
| *b606-b608* | $16.6 \pm 0.1$ | $0.60 \pm 0.36$ | $54.87 \pm 21.61$ | $53.96 \pm 19.06$ | $10.44 \pm 2.46$ | $10.34 \pm 2.06$ |
| *b606-b614* | $36.3 \pm 0.9$ | $0.11 \pm 0.16$ | $1.79 \pm 1.73$ | $1.89 \pm 1.49$ | $6.36 \pm 3.21$ | $6.78 \pm 2.85$ |

The overall B band *E*_EET_ of *a*612 to any other Chl *a* is 20/49% (FRET/TDC), arising mainly from *a*612-*a*611. For the reverse process (*a*611-*a*612) this is even higher, with an *E*_*EET*_ of 46/55%. Within the Chl *a*-*a* pairs, it can be seen that no other B band *a*-*a* processes are as efficient as *a*611-*a*612, with all other pairwise EET processes having an 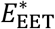 of 0.28 or less.

#### EET for B_x_ between Chls a--b

Chl *a*604 shows a similar picture for the B_x_ state compared to the Q_y_ state: Again, *a*604-*b*606 shows the strongest *V*_DA_, this time with 355.2/311.7 cm^−1^ (see Table 3), and rates of 20.3/15.3 ps^−1^. Despite *a*604-*b*607 pair being closer, *a*604-*b*608 displays the second strongest *V*_DA_ with 57.1/41.0 cm^−1^ due to the favorable orientation (κ^2^:1.41 compared to 0.07). 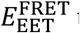 towards *b*606, *b*607, *b*608 and *b*614 are 0.26, 0.02, 0.01 and <0.01 and 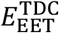 are <0.01 for all these transfers. This is due to preferential coupling to Crts, (see below).

#### EET for B_x_ between Chls b--b

For *V*_DA_ of Chl *b* with other Chl *b*, only the pair *b*606-*b*607 is significant. Contrary to the Q_y_ state, the *b*606-*b*607 *V*_DA_ is with 165.5/120.6 cm^−1^ in the same magnitude as the *a-a* and *a-b* couplings. While this *V*_DA_ is smaller than several *a*-*a* or *a*-*b* values, the number of possible *b*-*b* pairs (three) is too small to give an overview on general *b*-*b* coupling, compared to the other cases. It can be stated however that given favourable distance and orientation, EET is also possible between Chls *b*.

#### EET for B_x_ between Chls and Crts

Due to the strong absorbance of Crts in the Soret region, it is prudent to allow for Crts as EET acceptors. The B_x_ states of Chls indeed are found to interact with the Crts in a strong manner. Their couplings outcompete in the most cases all other processes. For example, *a*612-*L*620 shows *V*_DA_ of 1300.2/569.8 cm^−1^ (see Table 4) yielding rates of 83.7/15.4 ps^−1^ and *E*_EET_ of 64/23%. However, not every Chl has a Crt in the immediate vicinity. As an overview on the B band EET, Figure 3 shows the structural position and the *E*_EET_ of every Chl donor. The pie charts show the fractions of *E*_EET_.

**Table 4:** Comparison of the calculated average distances (*r*_DA_), κ^2^ values, Coulomb couplings of the FRET (*V*^FRET^) and TDC-scheme (*V*^TDC^) and the corresponding average FRET radii for the B_x_ state and the first excited state of the Crts (lutein (L), violaxanthin (V) and neoxanthin (N)).

| D-A Pair | $r_{DA}/\text{\AA}$ | $\kappa^2$ | $V^{\text{FRET}}/\text{cm}^{-1}$ | $V^{\text{TDC}}/\text{cm}^{-1}$ | $R_0^{\text{FRET}}/\text{\AA}$ | $R_0^{\text{TDC}}/\text{\AA}$ |
| --- | --- | --- | --- | --- | --- | --- |
| *a*612-*L*620 | 10.6±0.1 | 1.08±0.04 | 1300.22±368.65 | 569.79±83.53 | 11.86±0.66 | 9.11±0.23 |
| *a*612-*V*622 | 20.1±0.1 | 0.02±0.02 | 8.82±5.56 | 25.34±6.42 | 5.30±1.30 | 7.85±0.84 |
| *a*612-*N*623 | 28.5±0.4 | 0.08±0.14 | 5.39±5.77 | 4.29±3.15 | 7.22±5.32 | 7.07±2.03 |
| *a*604-*L*620 | 12.7±0.1 | 0.33±0.18 | 179.52±47.41 | 167.39±25.50 | 10.06±0.86 | 9.88±0.49 |
| *a*604-*V*622 | 17.4±0.1 | 0.17±0.08 | 50.30±15.01 | 69.65±37.21 | 9.29±0.90 | 10.07±2.14 |
| *a*604-*N*623 | 15.8±0.1 | 0.09±0.06 | 47.15±17.85 | 97.70±49.21 | 8.40±1.14 | 10.63±2.01 |
| *b*606-*L*620 | 18.8±0.1 | 0.88±0.06 | 112.73±16.79 | 84.12±13.26 | 11.69±0.86 | 10.60±0.79 |
| *b*606-*V*622 | 19.1±0.2 | 0.90±0.24 | 106.95±24.86 | 126.09±47.3 | 12.16±1.30 | 12.61±2.57 |
| *b*606-*N*623 | 12.1±0.2 | 2.26±0.16 | 642.04±97.57 | 367.06±60.79 | 14.49±1.06 | 12.02±0.86 |

**Figure 3:**
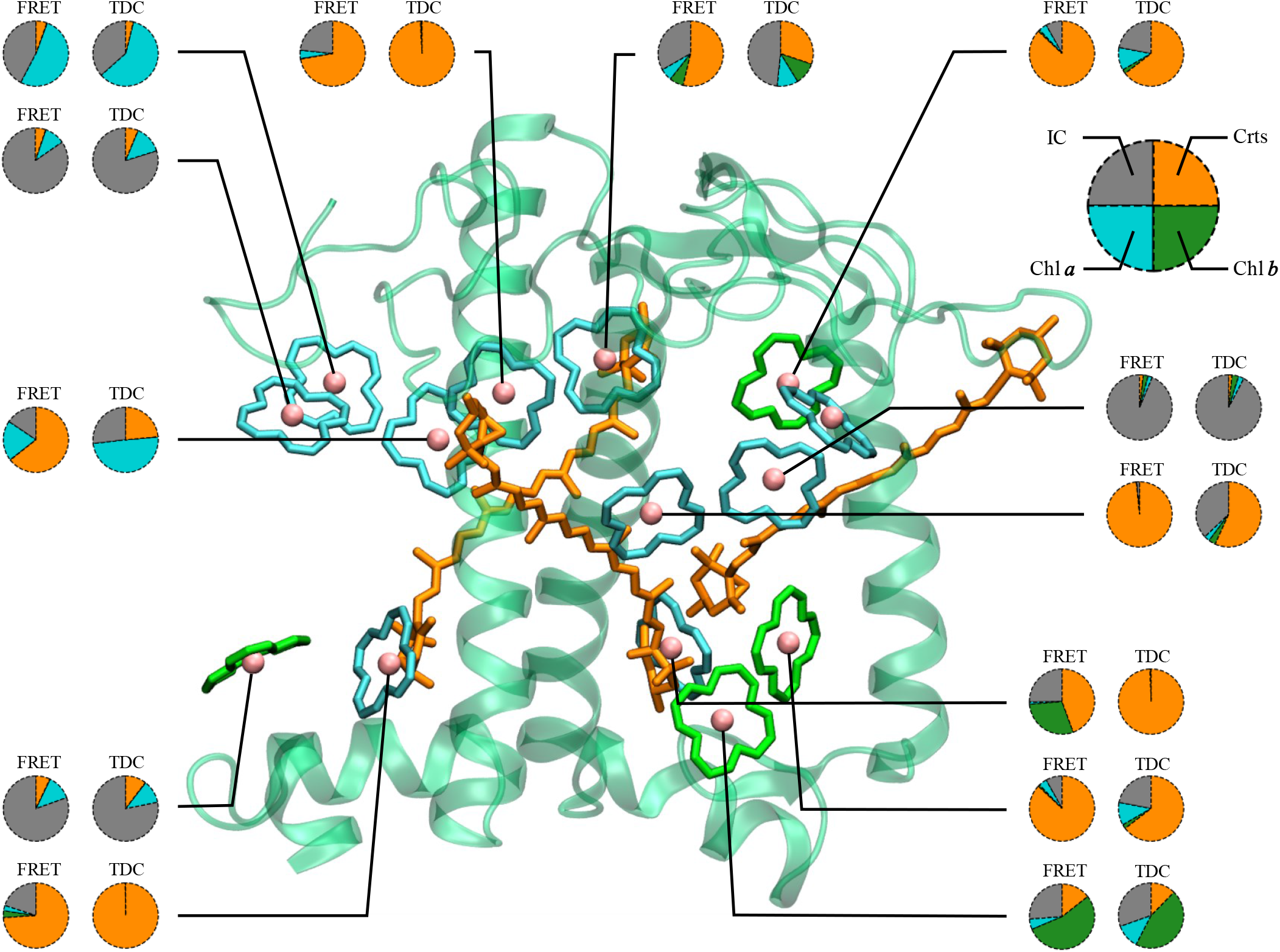
Structural representation of 13 Chls and efficiencies of IC processes (grey) and the E_EET_ of Bx states of a donor Chl towards a chromophores of type Chl a (cyan), Chl b (green), Crts (orange)).

**Figure 4:**
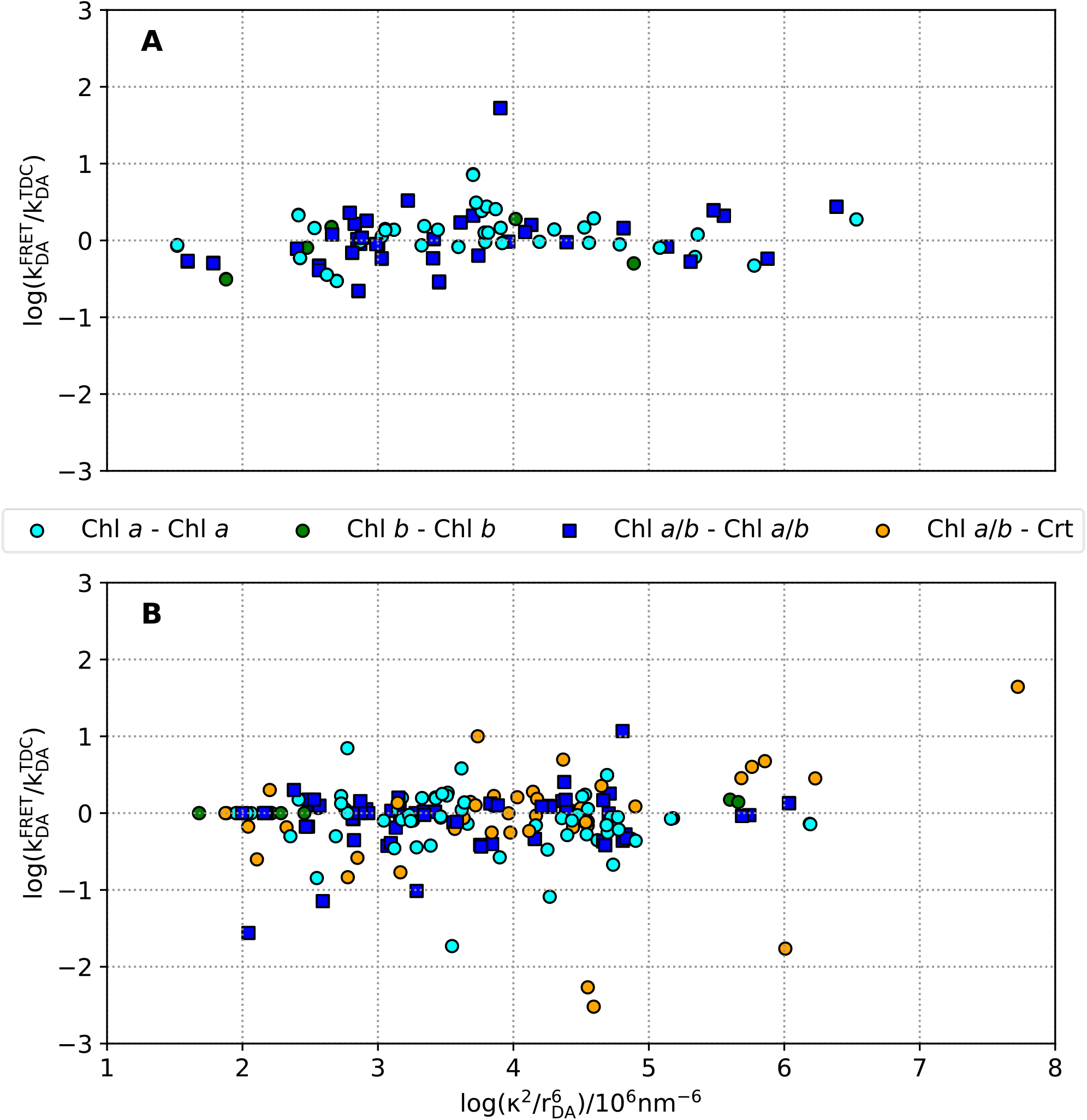
Ratio of FRET and TDC rates (log scale) depending on the FRET variables (κ^2^ and 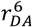) for Chl a-a couplings (cyan dots), b-b couplings (green dots), a-b couplings (blue squares) and Chl-Crt couplings (orange dots) of the Q_y_ state (A) and the B_x_ state (B).

**Figure 5:**
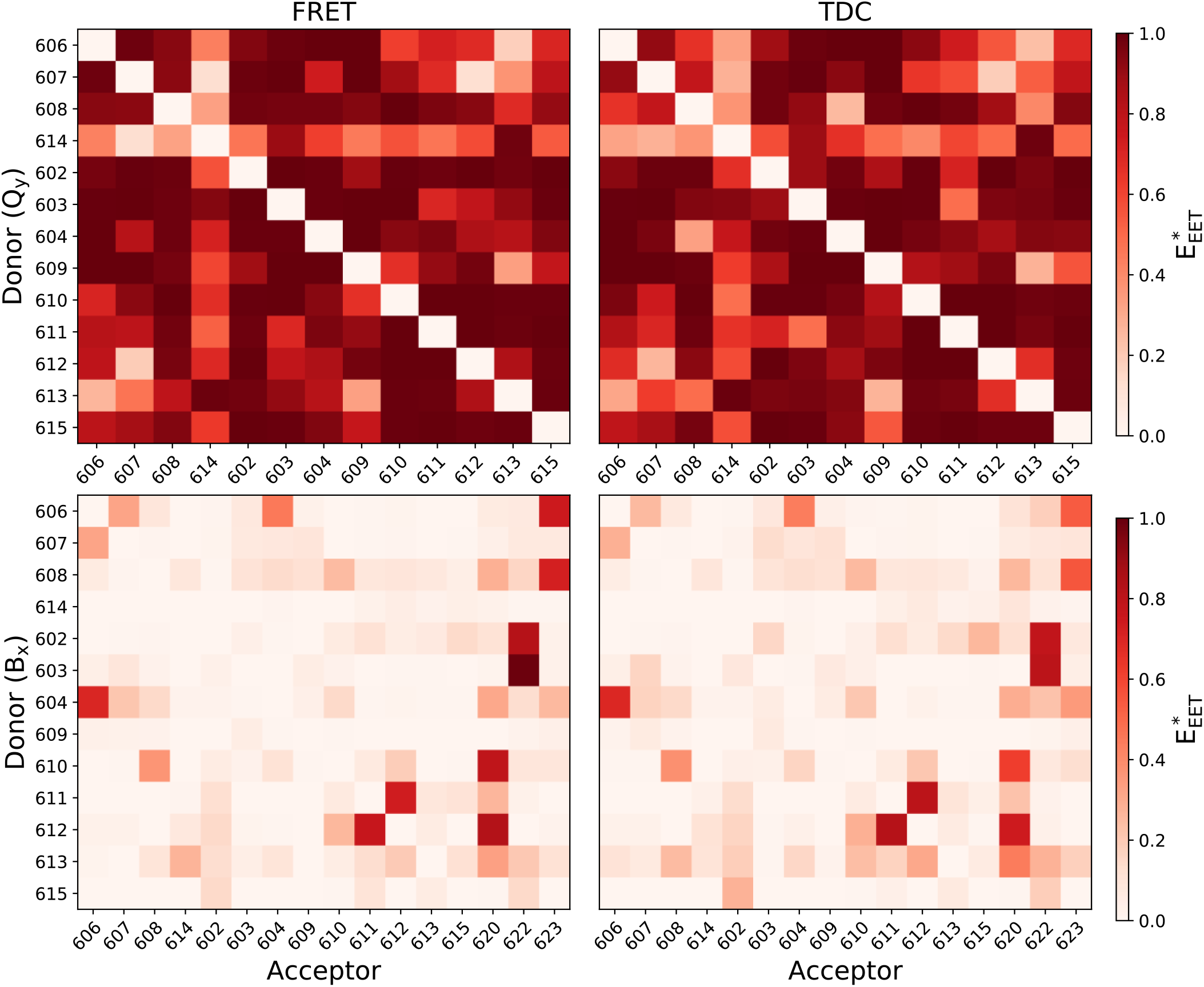
FRET (left) and TDC (right) 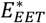 of the Q_y_ state (top) and the B_x_ state (bottom), while each row representing a Chl as donor and each column a Chl or Crt as acceptor.

The B band *E*_EET_ values are similar for FRET and TDC, apart from a few cases (see Table S13 and S14 in the SI). FRET and TDC show different pictures for *a*610 and *a*612. While FRET shows *E*_*EET*_ of 53% for *a*610-Crts and just 33% for IC processes, TDC shows *E*_*EET*_ of 30% for *a*610-Crts and 48% for IC processes. Chl *a*612 prefers the transfer to Crts with 0.64 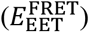 and shows 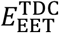 of 0.23 to Crts. In the most cases these differences are correlated with a single under- or overestimated coupling changing the overall *E*_EET_. As shown above, Chl *a*612 shows massively different *V*_DA_ values for the interaction *a*612-*L*620 for FRET and TDC. Without the strong coupling with the Lutein all other processes get more likely in TDC. The Chls *b*606, *b*608, *a*602, *a*603, *a*604 and *a*613 most likely transfer towards a Crt in both approaches. When neglecting the Crts as acceptors, the overall EET/IC ratio is almost independent of the FRET and TDC methods (⟨*E*_EET_⟩(Chl *a*/IC) = 0.55/0.64 and ⟨*E*_EET_⟩(Chl *b*/IC) = 0.45/0.33 for FRET/TDC in the B_x_ state). This means that between Chls, EET quality is well described by FRET, within about 0.1 accuracy compared to TDC.

Some Chls in our model may possibly show interactions which are not fully realistic in a PSII supercomplex arrangement. For example, Chl *b*614, *a*609 and *a*615 are preferring IC. All three Chls are peripheral and are close to the LHCII (*b*614 and *a*615) or to the PSII core complex (*a*609) or the omitted *a*616 of CP29, which forms the connection to the PSII core. As the model does not include surrounding complexes, some transfer partners may be missing. For all these cases, our results may underestimate the true EET.

Without doing explicit kinetics (prohibited due to the missing Crt emission), we can however already state that specific Chls indirectly transfer to Crts. For example, Chl *b*607 and *a*611 are strongly coupled with another Chl (*b*607-*b*606 and *a*611*-a*612) and therefor showing *E*_EET_ to Chl *b* (54/45%) and to Chl *a* (52/59%), respectively. However, Chl *b*606 itself is strongly coupled to the neoxanthin with *E*_EET_ of 0.58/0.38 (FRET/TDC) and Chl *a*612 is strongly coupled to the lutein with *E*_EET_ of 0.64/0.23 (FRET/TDC). Further Crt acceptor efficiency can be quantified by comparing a hypothetical Crt free system to the true CP29 arrangement. The most likely process that occurs in a Crt free system is IC as 8 of 13 Chls show an IC efficiency of above 0.50 for both approaches. Chl *b*606, *a*611 and *a*612 show *E*_EET_>0.50 to Chls *a* and Chl *b*607 show *E*_EET_>0.50 to Chls *b*. The Chl *a*604 differs slightly comparing the FRET and TDC values as the rates towards the Chls *b* are similar. While 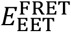 is with 0.53 higher than the IC efficiency with 0.45, the 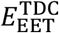 is with 0.46 lower than the IC efficiency with 0.50. Taking the Crts into account only Chl *b*614, *a*609 and *a*615 (and *a*610 in the TDC approach, which is described above) show IC as the most preferred process. Furthermore, *E*_EET_ of Chl *b*606-Crts is with 0.61/0.44 higher than the *E*_EET_ of Chl *b*606-Chls *a* with 0.20/0.28 (FRET/TDC).

In summary Figure 3 shows that most Chls are described in terms of *E*_EET_ the same way, regardless of FRET or TDC. Quantitatively, Chls *b*606, *b*607, *b*608, *b*614, *a*602, *a*609, *a*611, *a*613, *a*615 have virtually the same relative contribution in Figure 3, Chls *a*603, *a*604, *a*610 and *a*612 only differ slightly in terms of their EET to Crts.

### Q vs. B comparison

Figure 5 shows on overview on the pairwise Chl-Chl and Chl-Crt efficiencies for Q_y_ and B_x_ EET. It can be seen that the decay processes are not competitive for the Q_y_ state (top two plots), resulting in the well-understood high photosynthetic EET efficiency and quantum yield(cite). The bottom two plots of Figure 5 show how much this differs for the B_x_ state, where the decay processes are very competitive for many cases of the pairwise EET efficiency. However, most rows contain at least one Chl-Chl element and/or one Chl-Crt element of significant (>0.4) size. This indicates that EET is also possible here.

Looking at the individual coupling rates, FRET and TDC show a similar situation for the Q_y_ state, except for a single pair (*a*604*-b*608) with a factor of about 50 (see Figure 4A). As already discussed above, this outlier is a result of the PDA being less applicable for Chl *b*608, due to a coordination to a bridging water molecule. Figure 4B shows the difference of the rates calculated with the FRET and the TDC methods for the B_x_ state. Here, the transfer rates of the spatially more challenging Crts show the strongest differences for the FRET and TDC methods, with no clear trend upwards or downwards. This was expected, as TDC was specifically designed with these cases in mind. It is however interesting to see that the majority of Chl-Crt rates are still indifferent to the choice of method, only 5 Chl-Crt cases differ by an order of magnitude or more. Furthermore, Chl *a*613 differs in 5 cases for the B_x_ state by more than one order of magnitude. This is maybe correlated with the fact that the Chl *a*613 is close to Lut (4 out of 11 frames), which results in Lut becoming the Chl ligand. For those cases, both methods are likely unreliable.

In conclusion, FRET and TDC are mostly similar for the Q_y_ state and differ more strongly for the B_x_ state, especially taking the Crts as acceptors into account.

## Conclusion

In this article, we compared, for the CP29 complex, FRET and TDC regarding couplings, transfer rates, FRET radii (also analogously computed for TDC) and efficiencies. This analysis was performed both for the Q as well for the B band absorption region, the latter including Crt absorption. It was found that for the Q band, FRET or TDC treatment will not drastically affect the results, since the EET efficiencies within the network are dominated by coupling to the next neighbor and then almost always close to 1. For the B band, it was shown that EET between Chls in the B band exists and usually outpaces IC, except for those cases for which we omitted the likely acceptors in our model. The B band EET occurs regardless of coupling scheme and can have network efficiencies that are also close to unity, although not to individual Chl acceptors.

Our results show a similar picture for pure Chl-Chl coupling regardless of FRET or TDC and much more drastic changes when looking at Crt as acceptors. This is not surprising, since the TDC method was specifically set up to amend downsides of the FRET approach, namely the PDA. The PDA is less accurate for the spatially challenging Crts than for the Chls, thus we see the strongest changes in Crt acceptor efficiency (see Table 4) and only minor changes in the ratios of IC or Chl acceptor efficiencies.

As using TDC is not systematically affecting the interaction of two chromophores (consistently to higher or lower couplings), it is impossible to make a general statement regarding the impact of the multipole moments on a FRET result. In the investigated CP29 model, employing TDC indeed changes the coupling of Chls to Crts, and thus affects the resulting interaction between chromophores of an antenna complex like CP29. Which one represents the reality must be the task of future studies, but given that TDC is a better approximation to the Coulomb interaction between chromophores, it is likely that the TDC picture is closer to the real network.

The comparison of the Q and the B band shows how important the inclusion of the TDC method for analysis of the B_x_ state is. While for the Q_y_ states the decay processes are not competitive (each Chl shows at least a Γ_DA_ > 0.2 ps^−1^ to another Chl with a lifetime of *τ*_IC_ > 3.2 ns) this is not always the case for the B band. Here, much shorter lifetimes compete with similar or even higher couplings compared to the Q_y_ state. Yet, for many cases, EET remains the dominant B band process. Nevertheless, Chl *a*611 prefers a transfer band EET towards another Chl *a* and Chl *b*607 prefers a transfer band EET towards another Chl *b*. All other Chls are either close to a Crt, preferring this as target for their B band excitation energy or decay via IC processes. The actual strength of coupling to the Crts, depends on the coupling method (FRET or TDC).

The approximations used in our model (no Stokes shift, no spectral envelope and no Crt relaxation) unfortunately do not allow for stating explicit “blue” energy pathways, as opposed to “red” Q band pathways. However, we showed that the calculated model indicates that the Soret states are likely highly coupled, but curiously show almost no EET from Chl *a* to Chl *a*, despite Chl *a* being by far the most numerous chromophore in the system.

## Acknowledgements

The authors gratefully acknowledge funding by the Deutsche Forschungsgemeinschaft, project “Blue light pathways in light harvesting complexes”, project no. 393271229.

