## Supplementary material for "Impact of structural sampling, coupling scheme and state of interest on the energy transfer in CP29": Supporting_Information_TDC_final.pdf

List of Tables

Table 1: $Q_y$ FRET rates. 2
Table 2: $Q_y$ TDC rates. 3
Table 3: $Q_y$ FRET $E_{EET}^{*}$. 4
Table 4: $Q_y$ FRET $E_{EET}$. 5
Table 5: $Q_y$ TDC $E_{EET}^{*}$. 6
Table 6: $Q_y$ TDC $E_{EET}$. 7
Table 7: $B_x$ FRET rates. 8
Table 8: $B_x$ TDC rates. 9
Table 9: $B_x$ FRET $E_{EET}^{*}$. 10
Table 10: $B_x$ FRET $E_{EET}$. 11
Table 11: $B_x$ TDC $E_{EET}^{*}$. 12
Table 12: $B_x$ TDC $E_{EET}$. 13
Table 13: $B_x$ FRET $E_{EET}$. 14
Table 14: $B_x$ TDC $E_{EET}$. 14

List of Figures

Figure 1: Dynamical evolution of the $V_{\mathrm{FRET/TDC}}$, $R_0$ and $\kappa^2$ of Chl $a$612 Qy states to Chls $a$. 15
Figure 2: Dynamical evolution of the $V_{\mathrm{FRET/TDC}}$, $R_0$ and $\kappa^2$ of Chl a612 Bx states to Chls a.. 16

*Table 1: FRET rates in ps$^{-1}$ of $Q_y$ states for all chromophores with the italic letters indicating Chl a/b or Lutein (L), Violaxanthin (V) and Neoxanthin (N). Every row represents a donor and every column the respective acceptor.*

| | ***b*606** | ***b*607** | ***b*608** | ***b*614** | ***a*602** | ***a*603** | ***a*604** | ***a*609** | ***a*610** | ***a*611** | ***a*612** | ***a*613** | ***a*615** |
|---|---|---|---|---|---|---|---|---|---|---|---|---|---|
| ***b*606** | - | 0.05 | 0.01 | 0.00 | 0.01 | 0.03 | 3.04 | 0.18 | 0.00 | 0.00 | 0.00 | 0.00 | 0.00 |
| ***b*607** | 0.05 | - | 0.01 | 0.00 | 0.02 | 0.68 | 0.00 | 0.50 | 0.00 | 0.00 | 0.00 | 0.00 | 0.00 |
| ***b*608** | 0.01 | 0.01 | - | 0.00 | 0.01 | 0.02 | 0.01 | 0.07 | 0.97 | 0.01 | 0.01 | 0.00 | 0.00 |
| ***b*614** | 0.00 | 0.00 | 0.00 | - | 0.00 | 0.00 | 0.00 | 0.00 | 0.00 | 0.00 | 0.00 | 0.20 | 0.00 |
| ***a*602** | 0.01 | 0.02 | 0.01 | 0.00 | - | 0.11 | 0.02 | 0.02 | 0.04 | 0.01 | 0.09 | 0.01 | 1.62 |
| ***a*603** | 0.03 | 0.64 | 0.02 | 0.00 | 0.11 | - | 0.02 | 0.73 | 0.06 | 0.00 | 0.00 | 0.01 | 0.02 |
| ***a*604** | 2.85 | 0.00 | 0.01 | 0.00 | 0.02 | 0.02 | - | 0.10 | 0.01 | 0.00 | 0.00 | 0.00 | 0.00 |
| ***a*609** | 0.17 | 0.50 | 0.07 | 0.00 | 0.02 | 0.74 | 0.10 | - | 0.00 | 0.00 | 0.01 | 0.00 | 0.00 |
| ***a*610** | 0.00 | 0.00 | 0.91 | 0.00 | 0.04 | 0.06 | 0.01 | 0.00 | - | 0.36 | 0.62 | 0.02 | 0.02 |
| ***a*611** | 0.00 | 0.00 | 0.01 | 0.00 | 0.01 | 0.00 | 0.00 | 0.00 | 0.35 | - | 9.20 | 0.02 | 0.18 |
| ***a*612** | 0.00 | 0.00 | 0.00 | 0.00 | 0.09 | 0.00 | 0.00 | 0.01 | 0.63 | 9.54 | - | 0.00 | 0.02 |
| ***a*613** | 0.00 | 0.00 | 0.00 | 0.19 | 0.01 | 0.01 | 0.00 | 0.00 | 0.02 | 0.02 | 0.00 | - | 0.02 |
| ***a*615** | 0.00 | 0.00 | 0.00 | 0.00 | 1.54 | 0.02 | 0.00 | 0.00 | 0.02 | 0.17 | 0.01 | 0.02 | - |

*Table 2: TDC rates in ps*$^{-1}$ *of* $Q_y$ *states for all chromophores with the italic letters indicating Chl a/b or Lutein (L), Violaxanthin (V) and Neoxanthin (N). Every row represents a donor and every column the respective acceptor.*

| | ***b*606** | ***b*607** | ***b*608** | ***b*614** | ***a*602** | ***a*603** | ***a*604** | ***a*609** | ***a*610** | ***a*611** | ***a*612** | ***a*613** | ***a*615** |
|---|---|---|---|---|---|---|---|---|---|---|---|---|---|
| ***b*606** | - | 0.10 | 0.00 | 0.00 | 0.00 | 0.04 | 1.11 | 0.22 | 0.00 | 0.00 | 0.00 | 0.00 | 0.00 |
| ***b*607** | 0.10 | - | 0.00 | 0.00 | 0.02 | 0.32 | 0.01 | 0.20 | 0.00 | 0.00 | 0.00 | 0.00 | 0.00 |
| ***b*608** | 0.00 | 0.00 | - | 0.00 | 0.01 | 0.01 | 0.00 | 0.05 | 1.68 | 0.01 | 0.00 | 0.00 | 0.01 |
| ***b*614** | 0.00 | 0.00 | 0.00 | - | 0.00 | 0.00 | 0.00 | 0.00 | 0.00 | 0.00 | 0.00 | 0.38 | 0.00 |
| ***a*602** | 0.00 | 0.01 | 0.01 | 0.00 | - | 0.06 | 0.01 | 0.01 | 0.04 | 0.00 | 0.06 | 0.01 | 3.43 |
| ***a*603** | 0.03 | 0.31 | 0.01 | 0.00 | 0.06 | - | 0.02 | 1.21 | 0.04 | 0.00 | 0.00 | 0.01 | 0.02 |
| ***a*604** | 1.04 | 0.01 | 0.00 | 0.00 | 0.01 | 0.02 | - | 0.10 | 0.01 | 0.00 | 0.00 | 0.00 | 0.00 |
| ***a*609** | 0.21 | 0.20 | 0.05 | 0.00 | 0.01 | 1.22 | 0.10 | - | 0.00 | 0.00 | 0.00 | 0.00 | 0.00 |
| ***a*610** | 0.00 | 0.00 | 1.57 | 0.00 | 0.05 | 0.04 | 0.01 | 0.00 | - | 0.45 | 0.52 | 0.01 | 0.01 |
| ***a*611** | 0.00 | 0.00 | 0.01 | 0.00 | 0.00 | 0.00 | 0.00 | 0.00 | 0.44 | - | 4.90 | 0.01 | 0.20 |
| ***a*612** | 0.00 | 0.00 | 0.00 | 0.00 | 0.06 | 0.00 | 0.00 | 0.00 | 0.53 | 5.08 | - | 0.00 | 0.01 |
| ***a*613** | 0.00 | 0.00 | 0.00 | 0.36 | 0.01 | 0.02 | 0.00 | 0.00 | 0.01 | 0.01 | 0.00 | - | 0.02 |
| ***a*615** | 0.00 | 0.00 | 0.00 | 0.00 | 3.27 | 0.02 | 0.00 | 0.00 | 0.01 | 0.19 | 0.01 | 0.02 | - |

*Table 3: FRET $E^{*}_{EET}$ of $Q_y$ states for all chromophores with the italic letters indicating Chl a/b or Lutein (L), Violaxanthin (V) and Neoxanthin (N). Every row represents a donor and every column the respective acceptor. Values above 0.5 are highlighted bold characters.*

| | ***b*606** | ***b*607** | ***b*608** | ***b*614** | ***a*602** | ***a*603** | ***a*604** | ***a*609** | ***a*610** | ***a*611** | ***a*612** | ***a*613** | ***a*615** |
|---|---|---|---|---|---|---|---|---|---|---|---|---|---|
| ***b*606** | - | **0.98** | **0.93** | 0.43 | **0.94** | **0.99** | **>0.99** | **>0.99** | **0.62** | **0.72** | **0.68** | 0.18 | **0.70** |
| ***b*607** | **0.98** | - | **0.92** | 0.13 | **0.99** | **>0.99** | **0.74** | **>0.99** | **0.88** | **0.68** | 0.12 | 0.37 | **0.79** |
| ***b*608** | **0.93** | **0.92** | - | 0.33 | **0.97** | **0.97** | **0.97** | **0.94** | **>0.99** | **0.96** | **0.93** | **0.68** | **0.90** |
| ***b*614** | 0.43 | 0.13 | 0.33 | - | 0.46 | **0.89** | **0.62** | 0.45 | **0.56** | 0.46 | **0.58** | **0.98** | **0.54** |
| ***a*602** | **0.97** | **0.99** | **0.99** | **0.57** | - | **>0.99** | **0.99** | **0.88** | **>0.99** | **0.98** | **>0.99** | **0.97** | **>0.99** |
| ***a*603** | **0.99** | **>0.99** | **0.98** | **0.94** | **>0.99** | - | **0.99** | **>0.99** | **>0.99** | **0.70** | **0.78** | **0.91** | **0.99** |
| ***a*604** | **>0.99** | **0.82** | **0.98** | **0.72** | **0.99** | **0.99** | - | **>0.99** | **0.93** | **0.96** | **0.84** | **0.82** | **0.95** |
| ***a*609** | **>0.99** | **>0.99** | **0.97** | **0.60** | **0.88** | **>0.99** | **>0.99** | - | **0.67** | **0.91** | **0.97** | 0.33 | **0.78** |
| ***a*610** | **0.71** | **0.93** | **>0.99** | **0.67** | **>0.99** | **>0.99** | **0.93** | **0.66** | - | **>0.99** | **>0.99** | **0.99** | **0.99** |
| ***a*611** | **0.82** | **0.79** | **0.97** | **0.52** | **0.98** | **0.69** | **0.96** | **0.90** | **>0.99** | - | **>0.99** | **0.99** | **>0.99** |
| ***a*612** | **0.79** | 0.19 | **0.96** | **0.69** | **>0.99** | **0.78** | **0.84** | **0.97** | **>0.99** | **>0.99** | - | **0.84** | **0.99** |
| ***a*613** | 0.27 | 0.47 | **0.79** | **0.99** | **0.97** | **0.91** | **0.82** | 0.33 | **0.99** | **0.99** | **0.84** | - | **0.99** |
| ***a*615** | **0.80** | **0.87** | **0.94** | **0.63** | **>0.99** | **0.99** | **0.95** | **0.77** | **0.99** | **>0.99** | **0.98** | **0.99** | - |

*Table 4: FRET $E_{EET}$ of $Q_y$ states for all chromophores with the italic letters indicating Chl a/b or Lutein (L), Violaxanthin (V) and Neoxanthin (N). Every row represents a donor and every column the respective acceptor. The effective efficiency of the donor doing internal conversion (IC) is represented by the last column. Values above 0.5 are highlighted bold characters.*

| | ***b*606** | ***b*607** | ***b*608** | ***b*614** | ***a*602** | ***a*603** | ***a*604** | ***a*609** | ***a*610** | ***a*611** | ***a*612** | ***a*613** | ***a*615** | **IC** |
|---|---|---|---|---|---|---|---|---|---|---|---|---|---|---|
| ***b*606** | - | 0.02 | 0.00 | 0.00 | 0.00 | 0.01 | **0.91** | 0.06 | 0.00 | 0.00 | 0.00 | 0.00 | 0.00 | 0.00 |
| ***b*607** | 0.04 | - | 0.00 | 0.00 | 0.02 | **0.53** | 0.00 | 0.40 | 0.00 | 0.00 | 0.00 | 0.00 | 0.00 | 0.00 |
| ***b*608** | 0.01 | 0.00 | - | 0.00 | 0.01 | 0.02 | 0.01 | 0.06 | **0.88** | 0.01 | 0.00 | 0.00 | 0.00 | 0.00 |
| ***b*614** | 0.00 | 0.00 | 0.00 | - | 0.00 | 0.02 | 0.00 | 0.00 | 0.01 | 0.01 | 0.00 | **0.95** | 0.00 | 0.00 |
| ***a*602** | 0.00 | 0.01 | 0.01 | 0.00 | - | 0.06 | 0.01 | 0.01 | 0.02 | 0.01 | 0.05 | 0.00 | **0.82** | 0.00 |
| ***a*603** | 0.02 | 0.39 | 0.01 | 0.00 | 0.07 | - | 0.01 | 0.44 | 0.04 | 0.00 | 0.00 | 0.01 | 0.01 | 0.00 |
| ***a*604** | **0.94** | 0.00 | 0.00 | 0.00 | 0.01 | 0.01 | - | 0.03 | 0.00 | 0.00 | 0.00 | 0.00 | 0.00 | 0.00 |
| ***a*609** | 0.11 | 0.31 | 0.04 | 0.00 | 0.01 | 0.46 | 0.06 | - | 0.00 | 0.00 | 0.00 | 0.00 | 0.00 | 0.00 |
| ***a*610** | 0.00 | 0.00 | 0.44 | 0.00 | 0.02 | 0.03 | 0.00 | 0.00 | - | 0.18 | 0.30 | 0.01 | 0.01 | 0.00 |
| ***a*611** | 0.00 | 0.00 | 0.00 | 0.00 | 0.00 | 0.00 | 0.00 | 0.00 | 0.04 | - | **0.94** | 0.00 | 0.02 | 0.00 |
| ***a*612** | 0.00 | 0.00 | 0.00 | 0.00 | 0.01 | 0.00 | 0.00 | 0.00 | 0.06 | **0.93** | - | 0.00 | 0.00 | 0.00 |
| ***a*613** | 0.00 | 0.00 | 0.00 | **0.67** | 0.03 | 0.04 | 0.00 | 0.00 | 0.08 | 0.07 | 0.01 | - | 0.08 | 0.00 |
| ***a*615** | 0.00 | 0.00 | 0.00 | 0.00 | **0.86** | 0.01 | 0.00 | 0.00 | 0.01 | 0.10 | 0.01 | 0.01 | - | 0.00 |

*Table 5: TDC $E^*_{EET}$ of $Q_y$ states for all chromophores with the italic letters indicating Chl a/b or Lutein (L), Violaxanthin (V) and Neoxanthin (N). Every row represents a donor and every column the respective acceptor. Values above 0.5 are highlighted bold characters.*

| | ***b*606** | ***b*607** | ***b*608** | ***b*614** | ***a*602** | ***a*603** | ***a*604** | ***a*609** | ***a*610** | ***a*611** | ***a*612** | ***a*613** | ***a*615** |
|---|---|---|---|---|---|---|---|---|---|---|---|---|---|
| ***b*606** | - | **0.91** | **0.66** | 0.33 | **0.88** | **0.99** | **1.00** | **1.00** | **0.92** | **0.74** | **0.55** | 0.24 | **0.69** |
| ***b*607** | **0.91** | - | **0.78** | 0.28 | **0.97** | **0.99** | **0.93** | **1.00** | **0.64** | **0.58** | 0.19 | **0.53** | **0.78** |
| ***b*608** | **0.66** | **0.78** | - | 0.37 | **0.97** | **0.91** | 0.25 | **0.97** | **1.00** | **0.97** | **0.88** | 0.40 | **0.94** |
| ***b*614** | 0.32 | 0.28 | 0.37 | - | **0.58** | **0.89** | **0.66** | 0.48 | 0.41 | **0.60** | **0.50** | **0.98** | 0.49 |
| ***a*602** | **0.93** | **0.99** | **0.99** | **0.67** | - | **0.89** | **0.97** | **0.84** | **1.00** | **0.71** | **1.00** | **0.96** | **1.00** |
| ***a*603** | **0.99** | **1.00** | **0.94** | **0.94** | **0.89** | - | **0.99** | **1.00** | **1.00** | 0.49 | **0.95** | **0.96** | **0.99** |
| ***a*604** | **1.00** | **0.96** | **0.33** | **0.77** | **0.97** | **0.99** | - | **1.00** | **0.97** | **0.93** | **0.87** | **0.94** | **0.93** |
| ***a*609** | **1.00** | **1.00** | **0.99** | **0.62** | **0.84** | **1.00** | **1.00** | - | **0.83** | **0.88** | **0.95** | 0.27 | **0.56** |
| ***a*610** | **0.96** | **0.74** | **1.00** | 0.48 | **1.00** | **1.00** | **0.97** | **0.82** | - | **1.00** | **1.00** | **0.98** | **0.99** |
| ***a*611** | **0.83** | **0.70** | **0.98** | **0.65** | **0.71** | 0.48 | **0.93** | **0.88** | **1.00** | - | **1.00** | **0.96** | **1.00** |
| ***a*612** | **0.68** | 0.26 | **0.93** | **0.58** | **1.00** | **0.95** | **0.87** | **0.95** | **1.00** | **1.00** | - | **0.67** | **0.98** |
| ***a*613** | 0.31 | **0.62** | 0.49 | **0.99** | **0.96** | **0.96** | **0.94** | 0.27 | **0.98** | **0.96** | **0.67** | - | **0.99** |
| ***a*615** | **0.79** | **0.86** | **0.97** | **0.57** | **1.00** | **0.99** | **0.93** | **0.55** | **0.99** | **1.00** | **0.98** | **0.98** | - |

*Table 6: TDC $E_{EET}$ of $Q_y$ states for all chromophores with the italic letters indicating Chl a/b or Lutein (L), Violaxanthin (V) and Neoxanthin (N). Every row represents a donor and every column the respective acceptor. The effective efficiency of the donor doing internal conversion (IC) is represented by the last column. Values above 0.5 are highlighted bold characters.*

| | ***b*606** | ***b*607** | ***b*608** | ***b*614** | ***a*602** | ***a*603** | ***a*604** | ***a*609** | ***a*610** | ***a*611** | ***a*612** | ***a*613** | ***a*615** | **IC** |
|---|---|---|---|---|---|---|---|---|---|---|---|---|---|---|
| ***b*606** | - | 0.07 | 0.00 | 0.00 | 0.00 | 0.02 | **0.75** | 0.15 | 0.00 | 0.00 | 0.00 | 0.00 | 0.00 | 0.00 |
| ***b*607** | 0.15 | - | 0.01 | 0.00 | 0.02 | 0.49 | 0.02 | 0.31 | 0.00 | 0.00 | 0.00 | 0.00 | 0.00 | 0.00 |
| ***b*608** | 0.00 | 0.00 | - | 0.00 | 0.01 | 0.01 | 0.00 | 0.03 | **0.94** | 0.01 | 0.00 | 0.00 | 0.00 | 0.00 |
| ***b*614** | 0.00 | 0.00 | 0.00 | - | 0.00 | 0.01 | 0.00 | 0.00 | 0.00 | 0.01 | 0.00 | **0.97** | 0.00 | 0.00 |
| ***a*602** | 0.00 | 0.00 | 0.00 | 0.00 | - | 0.02 | 0.00 | 0.00 | 0.01 | 0.00 | 0.02 | 0.00 | **0.94** | 0.00 |
| ***a*603** | 0.02 | 0.18 | 0.01 | 0.00 | 0.03 | - | 0.01 | **0.70** | 0.03 | 0.00 | 0.00 | 0.01 | 0.01 | 0.00 |
| ***a*604** | **0.86** | 0.01 | 0.00 | 0.00 | 0.01 | 0.02 | - | 0.09 | 0.01 | 0.00 | 0.00 | 0.00 | 0.00 | 0.00 |
| ***a*609** | 0.12 | 0.11 | 0.03 | 0.00 | 0.00 | **0.68** | 0.06 | - | 0.00 | 0.00 | 0.00 | 0.00 | 0.00 | 0.00 |
| ***a*610** | 0.00 | 0.00 | 0.59 | 0.00 | 0.02 | 0.02 | 0.00 | 0.00 | - | 0.17 | 0.19 | 0.00 | 0.01 | 0.00 |
| ***a*611** | 0.00 | 0.00 | 0.00 | 0.00 | 0.00 | 0.00 | 0.00 | 0.00 | 0.08 | - | **0.88** | 0.00 | 0.04 | 0.00 |
| ***a*612** | 0.00 | 0.00 | 0.00 | 0.00 | 0.01 | 0.00 | 0.00 | 0.00 | 0.09 | **0.89** | - | 0.00 | 0.00 | 0.00 |
| ***a*613** | 0.00 | 0.00 | 0.00 | **0.86** | 0.01 | 0.04 | 0.01 | 0.00 | 0.02 | 0.02 | 0.01 | - | 0.04 | 0.00 |
| ***a*615** | 0.00 | 0.00 | 0.00 | 0.00 | **0.93** | 0.00 | 0.00 | 0.00 | 0.00 | 0.05 | 0.00 | 0.00 | - | 0.00 |

*Table 7: FRET rates in $ps^{-1}$ of $B_x$ states for all chromophores with the italic letters indicating Chl a/b or Lutein (L), Violaxanthin (V) and Neoxanthin (N). Every row represents a donor and every column the respective acceptor.*

| | ***b*606** | ***b*607** | ***b*608** | ***b*614** | ***a*602** | ***a*603** | ***a*604** | ***a*609** | ***a*610** | ***a*611** | ***a*612** | ***a*613** | ***a*615** | ***L*620** | ***V*622** | ***N*623** |
|---|---|---|---|---|---|---|---|---|---|---|---|---|---|---|---|---|
| ***b*606** | - | 10.70 | 0.67 | 0.00 | 0.01 | 0.23 | 17.18 | 0.14 | 0.01 | 0.01 | 0.02 | 0.00 | 0.01 | 1.00 | 1.26 | 51.39 |
| ***b*607** | 11.66 | - | 0.15 | 0.00 | 0.02 | 0.63 | 0.08 | 0.33 | 0.03 | 0.01 | 0.01 | 0.01 | 0.01 | 0.58 | 1.44 | 1.12 |
| ***b*608** | 0.49 | 0.12 | - | 0.01 | 0.03 | 0.08 | 0.23 | 0.15 | 2.34 | 0.00 | 0.00 | 0.02 | 0.00 | 4.27 | 1.28 | 56.09 |
| ***b*614** | 0.00 | 0.00 | 0.01 | - | 0.00 | 0.00 | 0.01 | 0.00 | 0.00 | 0.05 | 0.11 | 0.68 | 0.02 | 0.42 | 0.06 | 0.02 |
| ***a*602** | 0.02 | 0.02 | 0.06 | 0.01 | - | 0.21 | 0.03 | 0.02 | 0.36 | 0.74 | 0.31 | 0.43 | 1.00 | 0.70 | 52.42 | 0.19 |
| ***a*603** | 0.24 | 0.61 | 0.13 | 0.00 | 0.19 | - | 0.01 | 0.31 | 0.13 | 0.01 | 0.04 | 0.02 | 0.02 | 0.09 | $12e^3$ | 0.14 |
| ***a*604** | 17.62 | 1.56 | 0.82 | 0.04 | 0.04 | 0.02 | - | 0.04 | 0.54 | 0.01 | 0.07 | 0.06 | 0.00 | 8.78 | 1.76 | 19.86 |
| ***a*609** | 0.16 | 0.15 | 0.13 | 0.00 | 0.01 | 0.30 | 0.03 | - | 0.03 | 0.00 | 0.00 | 0.01 | 0.01 | 0.00 | 0.11 | 0.22 |
| ***a*610** | 0.01 | 0.03 | 3.57 | 0.00 | 0.32 | 0.13 | 0.70 | 0.03 | - | 0.41 | 1.51 | 0.02 | 0.03 | 26.74 | 0.55 | 0.56 |
| ***a*611** | 0.01 | 0.01 | 0.01 | 0.06 | 0.81 | 0.01 | 0.00 | 0.00 | 0.40 | - | 18.77 | 0.55 | 0.67 | 2.04 | 0.16 | 0.00 |
| ***a*612** | 0.03 | 0.01 | 0.00 | 0.13 | 0.34 | 0.03 | 0.02 | 0.00 | 1.45 | 19.61 | - | 0.68 | 0.01 | 71.31 | 0.01 | 0.00 |
| ***a*613** | 0.01 | 0.00 | 0.11 | 2.77 | 0.36 | 0.04 | 0.12 | 0.01 | 0.04 | 0.38 | 1.06 | - | 0.29 | 62.06 | 1.23 | 0.28 |
| ***a*615** | 0.01 | 0.01 | 0.00 | 0.02 | 1.00 | 0.02 | 0.00 | 0.00 | 0.03 | 0.66 | 0.02 | 0.32 | - | 0.03 | 1.02 | 0.00 |

*Table 8:TDC rates in $ps^{-1}$ of $B_x$ states for all chromophores with the italic letters indicating Chl a/b or Lutein (L), Violaxanthin (V) and Neoxanthin (N). Every row represents a donor and every column the respective acceptor.*

| | ***b*606** | ***b*607** | ***b*608** | ***b*614** | ***a*602** | ***a*603** | ***a*604** | ***a*609** | ***a*610** | ***a*611** | ***a*612** | ***a*613** | ***a*615** | ***L*620** | ***V*622** | ***N*623** |
|---|---|---|---|---|---|---|---|---|---|---|---|---|---|---|---|---|
| ***b*606** | - | 7.14 | 0.63 | 0.00 | 0.01 | 0.22 | 12.71 | 0.10 | 0.02 | 0.01 | 0.02 | 0.08 | 0.00 | 0.62 | 2.26 | 18.03 |
| ***b*607** | 8.37 | - | 0.12 | 0.00 | 0.02 | 1.19 | 0.20 | 0.76 | 0.04 | 0.01 | 0.01 | 0.02 | 0.01 | 0.35 | 0.76 | 1.21 |
| ***b*608** | 0.45 | 0.09 | - | 0.01 | 0.02 | 0.07 | 0.13 | 0.10 | 2.49 | 0.00 | 0.00 | 0.11 | 0.00 | 1.89 | 0.83 | 14.01 |
| ***b*614** | 0.00 | 0.00 | 0.01 | - | 0.01 | 0.00 | 0.02 | 0.00 | 0.00 | 0.14 | 0.23 | 0.41 | 0.02 | 0.68 | 0.05 | 0.03 |
| ***a*602** | 0.01 | 0.02 | 0.05 | 0.01 | - | 2.56 | 0.09 | 0.01 | 0.21 | 0.83 | 0.38 | 0.96 | 2.15 | 1.01 | $3e^3$ | 1.11 |
| ***a*603** | 0.22 | 1.28 | 0.11 | 0.00 | 0.56 | - | 0.04 | 0.44 | 0.19 | 0.01 | 0.02 | 0.01 | 0.02 | 0.01 | 26.18 | 0.25 |
| ***a*604** | 13.13 | 0.62 | 1.91 | 0.02 | 0.03 | 0.05 | - | 0.06 | 0.97 | 0.00 | 0.04 | 0.14 | 0.00 | 1.77 | 2.03 | $7e^3$ |
| ***a*609** | 0.11 | 0.40 | 0.11 | 0.00 | 0.01 | 0.44 | 0.03 | - | 0.02 | 0.00 | 0.00 | 0.01 | 0.00 | 0.00 | 0.08 | 0.22 |
| ***a*610** | 0.03 | 0.04 | 3.88 | 0.00 | 0.19 | 0.19 | 1.14 | 0.02 | - | 0.36 | 1.76 | 0.02 | 0.03 | 9.44 | 0.47 | 0.85 |
| ***a*611** | 0.01 | 0.01 | 0.00 | 0.17 | 0.91 | 0.01 | 0.00 | 0.00 | 0.36 | - | 25.88 | 0.68 | 0.21 | 1.67 | 0.13 | 0.01 |
| ***a*612** | 0.03 | 0.01 | 0.00 | 0.27 | 0.43 | 0.02 | 0.04 | 0.00 | 1.72 | 27.32 | - | 2.25 | 0.02 | 15.03 | 0.04 | 0.00 |
| ***a*613** | 0.22 | 0.06 | 1.09 | 0.24 | 0.68 | 0.01 | 0.46 | 0.01 | 2.21 | 0.72 | 4.95 | - | 0.33 | $12e^3$ | 2.10 | 1.08 |
| ***a*615** | 0.00 | 0.01 | 0.00 | 0.02 | 2.30 | 0.02 | 0.00 | 0.00 | 0.03 | 0.21 | 0.02 | 0.40 | - | 0.03 | 1.33 | 0.00 |

*Table 9: FRET $E^{*}_{EET}$ of $B_x$ states for all chromophores with the italic letters indicating Chl a/b or Lutein (L), Violaxanthin (V) and Neoxanthin (N). Every row represents a donor and every column the respective acceptor. Values above 0.5 are highlighted bold characters.*

| | ***b*606** | ***b*607** | ***b*608** | ***b*614** | ***a*602** | ***a*603** | ***a*604** | ***a*609** | ***a*610** | ***a*611** | ***a*612** | ***a*613** | ***a*615** | ***L*620** | ***V*622** | ***N*623** |
|---|---|---|---|---|---|---|---|---|---|---|---|---|---|---|---|---|
| ***b*606** | - | 0.32 | 0.09 | 0.00 | 0.01 | 0.08 | 0.45 | 0.03 | 0.00 | 0.01 | 0.01 | 0.01 | 0.00 | 0.06 | 0.07 | **0.75** |
| ***b*607** | 0.32 | - | 0.01 | 0.00 | 0.01 | 0.07 | 0.08 | 0.09 | 0.00 | 0.01 | 0.01 | 0.00 | 0.01 | 0.03 | 0.07 | 0.07 |
| ***b*608** | 0.06 | 0.01 | - | 0.08 | 0.01 | 0.10 | 0.14 | 0.12 | 0.25 | 0.08 | 0.09 | 0.11 | 0.04 | 0.28 | 0.16 | **0.72** |
| ***b*614** | 0.00 | 0.00 | 0.00 | - | 0.00 | 0.00 | 0.01 | 0.00 | 0.00 | 0.03 | 0.05 | 0.11 | 0.04 | 0.02 | 0.00 | 0.01 |
| ***a*602** | 0.00 | 0.00 | 0.01 | 0.00 | - | 0.04 | 0.01 | 0.00 | 0.06 | 0.11 | 0.05 | 0.07 | 0.14 | 0.11 | **0.82** | 0.03 |
| ***a*603** | 0.04 | 0.09 | 0.02 | 0.00 | 0.03 | - | 0.00 | 0.05 | 0.02 | 0.00 | 0.01 | 0.00 | 0.00 | 0.02 | **0.99** | 0.02 |
| ***a*604** | **0.70** | 0.21 | 0.15 | 0.02 | 0.02 | 0.01 | - | 0.04 | 0.15 | 0.00 | 0.01 | 0.07 | 0.00 | 0.31 | 0.13 | 0.26 |
| ***a*609** | 0.03 | 0.03 | 0.02 | 0.00 | 0.00 | 0.05 | 0.01 | - | 0.01 | 0.00 | 0.00 | 0.00 | 0.00 | 0.00 | 0.02 | 0.03 |
| ***a*610** | 0.00 | 0.01 | 0.36 | 0.00 | 0.05 | 0.02 | 0.11 | 0.01 | - | 0.07 | 0.19 | 0.00 | 0.01 | **0.79** | 0.09 | 0.09 |
| ***a*611** | 0.00 | 0.00 | 0.00 | 0.01 | 0.12 | 0.00 | 0.00 | 0.00 | 0.07 | - | **0.73** | 0.08 | 0.10 | 0.26 | 0.03 | 0.00 |
| ***a*612** | 0.03 | 0.02 | 0.00 | 0.08 | 0.15 | 0.01 | 0.01 | 0.00 | 0.26 | **0.76** | - | 0.15 | 0.00 | **0.83** | 0.00 | 0.02 |
| ***a*613** | 0.01 | 0.00 | 0.10 | 0.27 | 0.13 | 0.04 | 0.09 | 0.00 | 0.05 | 0.13 | 0.20 | - | 0.11 | 0.33 | 0.21 | 0.11 |
| ***a*615** | 0.00 | 0.00 | 0.00 | 0.00 | 0.15 | 0.00 | 0.00 | 0.00 | 0.01 | 0.10 | 0.00 | 0.05 | - | 0.01 | 0.15 | 0.00 |

*Table 10: FRET $E_{EET}$ of $B_x$ states for all chromophores with the italic letters indicating Chl a/b or Lutein (L), Violaxanthin (V) and Neoxanthin (N). Every row represents a donor and every column the respective acceptor. The effective efficiency of the donor doing internal conversion (IC) is represented by the last column. Values above 0.5 are highlighted bold characters.*

| | ***b*606** | ***b*607** | ***b*608** | ***b*614** | ***a*602** | ***a*603** | ***a*604** | ***a*609** | ***a*610** | ***a*611** | ***a*612** | ***a*613** | ***a*615** | ***L*620** | ***V*622** | ***N*623** | ***IC*** |
|---|---|---|---|---|---|---|---|---|---|---|---|---|---|---|---|---|---|
| ***b*606** | - | 0.12 | 0.01 | 0.00 | 0.00 | 0.00 | 0.19 | 0.00 | 0.00 | 0.00 | 0.00 | 0.00 | 0.00 | 0.01 | 0.01 | **0.58** | 0.06 |
| ***b*607** | **0.54** | - | 0.01 | 0.00 | 0.00 | 0.03 | 0.00 | 0.02 | 0.00 | 0.00 | 0.00 | 0.00 | 0.00 | 0.03 | 0.07 | 0.05 | 0.26 |
| ***b*608** | 0.01 | 0.00 | - | 0.00 | 0.00 | 0.00 | 0.00 | 0.00 | 0.03 | 0.00 | 0.00 | 0.00 | 0.00 | 0.06 | 0.02 | **0.79** | 0.08 |
| ***b*614** | 0.00 | 0.00 | 0.00 | - | 0.00 | 0.00 | 0.00 | 0.00 | 0.00 | 0.01 | 0.01 | 0.10 | 0.00 | 0.06 | 0.01 | 0.00 | **0.80** |
| ***a*602** | 0.00 | 0.00 | 0.00 | 0.00 | - | 0.00 | 0.00 | 0.00 | 0.00 | 0.01 | 0.00 | 0.01 | 0.01 | 0.01 | **0.71** | 0.00 | 0.23 |
| ***a*603** | 0.00 | 0.00 | 0.00 | 0.00 | 0.00 | - | 0.00 | 0.00 | 0.00 | 0.00 | 0.00 | 0.00 | 0.00 | 0.00 | **0.98** | 0.00 | 0.01 |
| ***a*604** | 0.26 | 0.02 | 0.01 | 0.00 | 0.00 | 0.00 | - | 0.00 | 0.01 | 0.00 | 0.00 | 0.00 | 0.00 | 0.13 | 0.03 | 0.29 | 0.25 |
| ***a*609** | 0.01 | 0.01 | 0.01 | 0.00 | 0.00 | 0.02 | 0.00 | - | 0.00 | 0.00 | 0.00 | 0.00 | 0.00 | 0.00 | 0.01 | 0.01 | **0.94** |
| ***a*610** | 0.00 | 0.00 | 0.07 | 0.00 | 0.01 | 0.00 | 0.01 | 0.00 | - | 0.01 | 0.03 | 0.00 | 0.00 | **0.52** | 0.01 | 0.01 | 0.33 |
| ***a*611** | 0.00 | 0.00 | 0.00 | 0.00 | 0.02 | 0.00 | 0.00 | 0.00 | 0.01 | - | 0.46 | 0.01 | 0.02 | 0.05 | 0.00 | 0.00 | 0.42 |
| ***a*612** | 0.00 | 0.00 | 0.00 | 0.00 | 0.00 | 0.00 | 0.00 | 0.00 | 0.01 | 0.18 | - | 0.01 | 0.00 | **0.64** | 0.00 | 0.00 | 0.16 |
| ***a*613** | 0.00 | 0.00 | 0.00 | 0.03 | 0.00 | 0.00 | 0.00 | 0.00 | 0.00 | 0.00 | 0.01 | - | 0.00 | **0.72** | 0.01 | 0.00 | 0.20 |
| ***a*615** | 0.00 | 0.00 | 0.00 | 0.00 | 0.05 | 0.00 | 0.00 | 0.00 | 0.00 | 0.03 | 0.00 | 0.02 | - | 0.00 | 0.05 | 0.00 | **0.85** |

*Table 11: TDC $E^{*}_{EET}$ of $B_x$ states for all chromophores with the italic letters indicating Chl a/b or Lutein (L), Violaxanthin (V) and Neoxanthin (N). Every row represents a donor and every column the respective acceptor. Values above 0.5 are highlighted bold characters.*

| | ***b*606** | ***b*607** | ***b*608** | ***b*614** | ***a*602** | ***a*603** | ***a*604** | ***a*609** | ***a*610** | ***a*611** | ***a*612** | ***a*613** | ***a*615** | ***L*620** | ***V*622** | ***N*623** |
|---|---|---|---|---|---|---|---|---|---|---|---|---|---|---|---|---|
| ***b*606** | - | 0.25 | 0.07 | 0.00 | 0.01 | 0.08 | 0.44 | 0.03 | 0.01 | 0.01 | 0.01 | 0.08 | 0.00 | 0.10 | 0.18 | **0.54** |
| ***b*607** | 0.28 | - | 0.01 | 0.00 | 0.02 | 0.14 | 0.09 | 0.12 | 0.00 | 0.01 | 0.01 | 0.04 | 0.00 | 0.06 | 0.08 | 0.09 |
| ***b*608** | 0.05 | 0.01 | - | 0.09 | 0.00 | 0.10 | 0.13 | 0.11 | 0.25 | 0.08 | 0.09 | 0.15 | 0.02 | 0.26 | 0.11 | **0.55** |
| ***b*614** | 0.00 | 0.00 | 0.00 | - | 0.00 | 0.00 | 0.01 | 0.00 | 0.00 | 0.04 | 0.07 | 0.08 | 0.03 | 0.09 | 0.02 | 0.01 |
| ***a*602** | 0.00 | 0.00 | 0.01 | 0.00 | - | 0.16 | 0.02 | 0.00 | 0.04 | 0.12 | 0.06 | 0.14 | 0.26 | 0.12 | **0.78** | 0.08 |
| ***a*603** | 0.04 | 0.16 | 0.02 | 0.00 | 0.08 | - | 0.01 | 0.07 | 0.03 | 0.00 | 0.00 | 0.00 | 0.00 | 0.00 | **0.80** | 0.04 |
| ***a*604** | **0.69** | 0.17 | 0.15 | 0.02 | 0.02 | 0.05 | - | 0.05 | 0.21 | 0.00 | 0.03 | 0.09 | 0.01 | 0.29 | 0.22 | 0.35 |
| ***a*609** | 0.02 | 0.06 | 0.02 | 0.00 | 0.00 | 0.07 | 0.01 | - | 0.00 | 0.00 | 0.00 | 0.00 | 0.00 | 0.00 | 0.01 | 0.04 |
| ***a*610** | 0.01 | 0.01 | 0.38 | 0.00 | 0.03 | 0.03 | 0.16 | 0.00 | - | 0.06 | 0.21 | 0.00 | 0.01 | **0.62** | 0.08 | 0.13 |
| ***a*611** | 0.00 | 0.00 | 0.00 | 0.03 | 0.14 | 0.00 | 0.00 | 0.00 | 0.06 | - | **0.80** | 0.10 | 0.04 | 0.23 | 0.02 | 0.00 |
| ***a*612** | 0.03 | 0.02 | 0.00 | 0.11 | 0.16 | 0.01 | 0.03 | 0.00 | 0.28 | **0.82** | - | 0.15 | 0.01 | **0.74** | 0.03 | 0.00 |
| ***a*613** | 0.10 | 0.07 | 0.25 | 0.10 | 0.19 | 0.00 | 0.15 | 0.02 | 0.24 | 0.17 | 0.31 | - | 0.06 | 0.44 | 0.28 | 0.18 |
| ***a*615** | 0.00 | 0.00 | 0.00 | 0.00 | 0.28 | 0.00 | 0.00 | 0.00 | 0.01 | 0.04 | 0.00 | 0.06 | - | 0.00 | 0.18 | 0.00 |

*Table 12: TDC $E_{EET}$ of $B_x$ states for all chromophores with the italic letters indicating Chl a/b or Lutein (L). Violaxanthin (V) and Neoxanthin (N). Every row represents a donor and every column the respective acceptor. The effective efficiency of the donor doing internal conversion (IC) is represented by the last column.*

| | ***b*606** | ***b*607** | ***b*608** | ***b*614** | ***a*602** | ***a*603** | ***a*604** | ***a*609** | ***a*610** | ***a*611** | ***a*612** | ***a*613** | ***a*615** | ***L*620** | ***V*622** | ***N*623** | ***IC*** |
|---|---|---|---|---|---|---|---|---|---|---|---|---|---|---|---|---|---|
| ***b*606** | - | 0.15 | 0.01 | 0.00 | 0.00 | 0.00 | 0.27 | 0.00 | 0.00 | 0.00 | 0.00 | 0.00 | 0.00 | 0.01 | 0.05 | 0.38 | 0.12 |
| ***b*607** | 0.45 | - | 0.01 | 0.00 | 0.00 | 0.06 | 0.01 | 0.04 | 0.00 | 0.00 | 0.00 | 0.00 | 0.00 | 0.02 | 0.04 | 0.06 | 0.30 |
| ***b*608** | 0.02 | 0.00 | - | 0.00 | 0.00 | 0.00 | 0.00 | 0.00 | 0.10 | 0.00 | 0.00 | 0.00 | 0.00 | 0.07 | 0.03 | **0.54** | 0.22 |
| ***b*614** | 0.00 | 0.00 | 0.00 | - | 0.00 | 0.00 | 0.00 | 0.00 | 0.00 | 0.02 | 0.03 | 0.06 | 0.00 | 0.09 | 0.01 | 0.00 | **0.78** |
| ***a*602** | 0.00 | 0.00 | 0.00 | 0.00 | - | 0.00 | 0.00 | 0.00 | 0.00 | 0.00 | 0.00 | 0.00 | 0.00 | 0.00 | **0.99** | 0.00 | 0.01 |
| ***a*603** | 0.00 | 0.03 | 0.00 | 0.00 | 0.01 | - | 0.00 | 0.01 | 0.00 | 0.00 | 0.00 | 0.00 | 0.00 | 0.00 | **0.56** | 0.01 | 0.37 |
| ***a*604** | 0.00 | 0.00 | 0.00 | 0.00 | 0.00 | 0.00 | - | 0.00 | 0.00 | 0.00 | 0.00 | 0.00 | 0.00 | 0.00 | 0.00 | **0.99** | 0.00 |
| ***a*609** | 0.01 | 0.02 | 0.01 | 0.00 | 0.00 | 0.02 | 0.00 | - | 0.00 | 0.00 | 0.00 | 0.00 | 0.00 | 0.00 | 0.00 | 0.01 | **0.92** |
| ***a*610** | 0.00 | 0.00 | 0.11 | 0.00 | 0.01 | 0.01 | 0.03 | 0.00 | - | 0.01 | 0.05 | 0.00 | 0.00 | 0.26 | 0.01 | 0.02 | 0.48 |
| ***a*611** | 0.00 | 0.00 | 0.00 | 0.00 | 0.02 | 0.00 | 0.00 | 0.00 | 0.01 | - | **0.55** | 0.01 | 0.00 | 0.04 | 0.00 | 0.00 | 0.36 |
| ***a*612** | 0.00 | 0.00 | 0.00 | 0.00 | 0.01 | 0.00 | 0.00 | 0.00 | 0.03 | 0.42 | - | 0.03 | 0.00 | 0.23 | 0.00 | 0.00 | 0.27 |
| ***a*613** | 0.00 | 0.00 | 0.00 | 0.00 | 0.00 | 0.00 | 0.00 | 0.00 | 0.00 | 0.00 | 0.00 | - | 0.00 | **1.00** | 0.00 | 0.00 | 0.00 |
| ***a*615** | 0.00 | 0.00 | 0.00 | 0.00 | 0.11 | 0.00 | 0.00 | 0.00 | 0.00 | 0.01 | 0.00 | 0.02 | - | 0.00 | 0.06 | 0.00 | **0.80** |

*Table 13: FRET $E_{EET}$ in % of $B_x$ states for all Chl towards a type of chromophores (e.g. Chl b606-Chls b: 12.9%). Every row represents a donor and every column the respective type of chromophores (Crts including Lutein. Violaxanthin. Neoxanthin). The effective efficiency of the donor doing internal conversion (IC) is represented by the last column.*

| | Chl *b* | Chl *a* | Crts | IC |
|---|---|---|---|---|
| *b*606 | 12.9 | 19.9 | 60.7 | 6.5 |
| *b*607 | 54.2 | 5.2 | 14.4 | 26.2 |
| *b*608 | 0.9 | 4.0 | 87.0 | 8.1 |
| *b*614 | 0.2 | 12.4 | 7.0 | 80.5 |
| *a*602 | 0.1 | 4.2 | 72.3 | 23.4 |
| *a*603 | 0.1 | 0.1 | 98.4 | 1.5 |
| *a*604 | 29.3 | 1.1 | 44.4 | 25.2 |
| *a*609 | 2.4 | 2.1 | 1.8 | 93.7 |
| *a*610 | 7.0 | 6.1 | 53.7 | 33.3 |
| *a*611 | 0.2 | 52.0 | 5.4 | 42.3 |
| *a*612 | 0.2 | 20.0 | 64.3 | 15.6 |
| *a*613 | 3.4 | 2.7 | 73.9 | 20.1 |
| *a*615 | 0.2 | 10.1 | 5.2 | 84.6 |

*Table 14: TDC $E_{EET}$ in % of $B_x$ states for all Chl towards a type of chromophores (e.g. Chl b606-Chls b: 16.3%). Every row represents a donor and every column the respective type of chromophores (Crts including Lutein. Violaxanthin. Neoxanthin). The effective efficiency of the donor doing internal conversion (IC) is represented by the last column.*

| | Chl *b* | Chl *a* | Crts | IC |
|---|---|---|---|---|
| *b*606 | 16.3 | 27.7 | 43.9 | 12.0 |
| *b*607 | 45.2 | 12.0 | 12.3 | 30.4 |
| *b*608 | 2.1 | 11.3 | 64.6 | 22.0 |
| *b*614 | 0.1 | 11.4 | 10.4 | 78.1 |
| *a*602 | 0.0 | 0.2 | 99.2 | 0.6 |
| *a*603 | 3.4 | 2.8 | 56.8 | 37.0 |
| *a*604 | 0.2 | 0.0 | 99.5 | 0.3 |
| *a*609 | 3.3 | 2.7 | 1.6 | 92.3 |
| *a*610 | 11.1 | 10.4 | 30.2 | 48.3 |
| *a*611 | 0.4 | 59.3 | 3.8 | 36.5 |
| *a*612 | 0.5 | 49.4 | 23.4 | 26.8 |
| *a*613 | 0.0 | 0.1 | 99.8 | 0.1 |
| *a*615 | 0.1 | 13.8 | 6.3 | 79.8 |

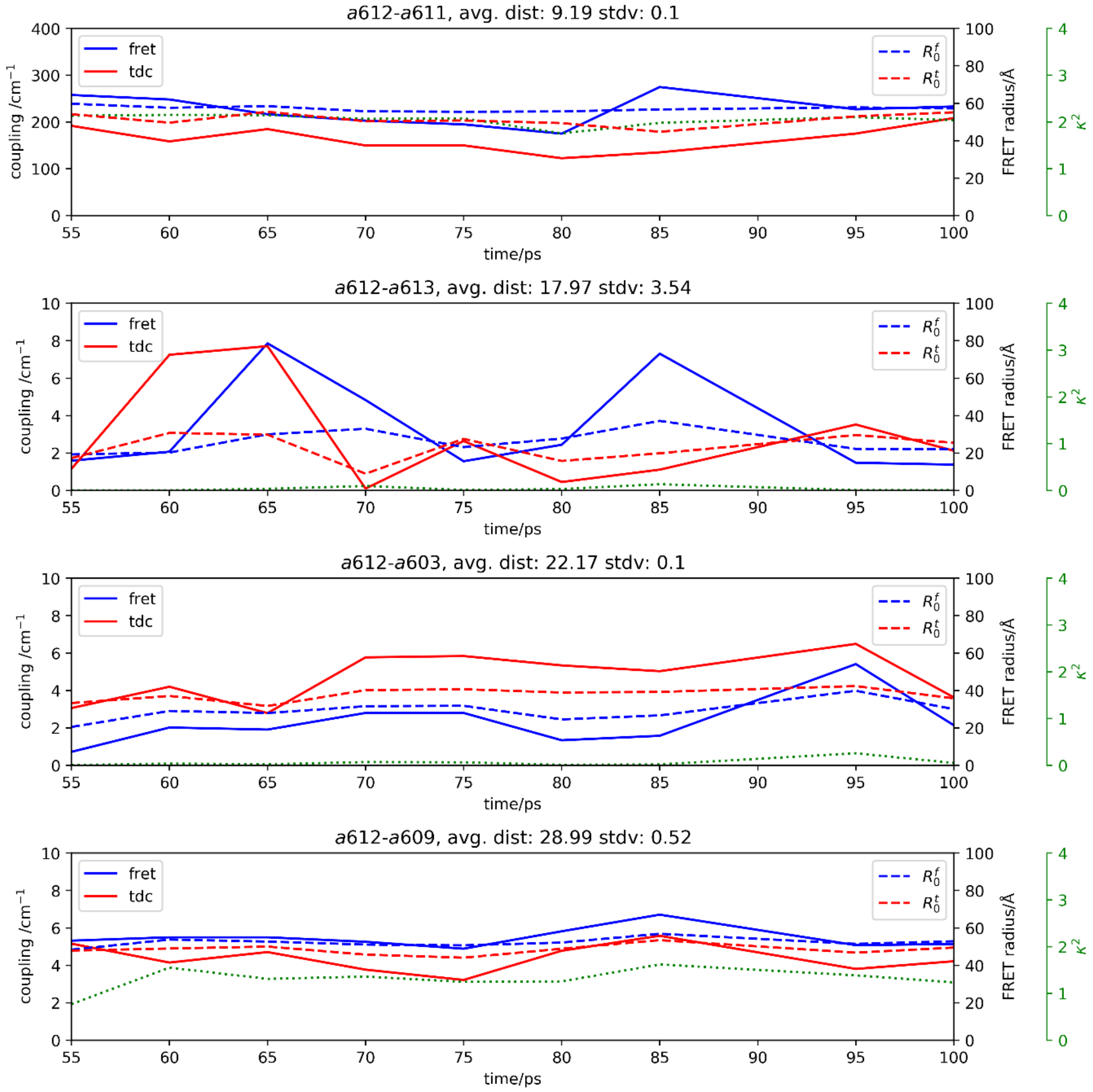


*Figure 1: $Q_y$ couplings from the FRET and TDC methods and the corresponding FRET radii and $\kappa^2$ values, between different Chl a-a pairs (A-D).*

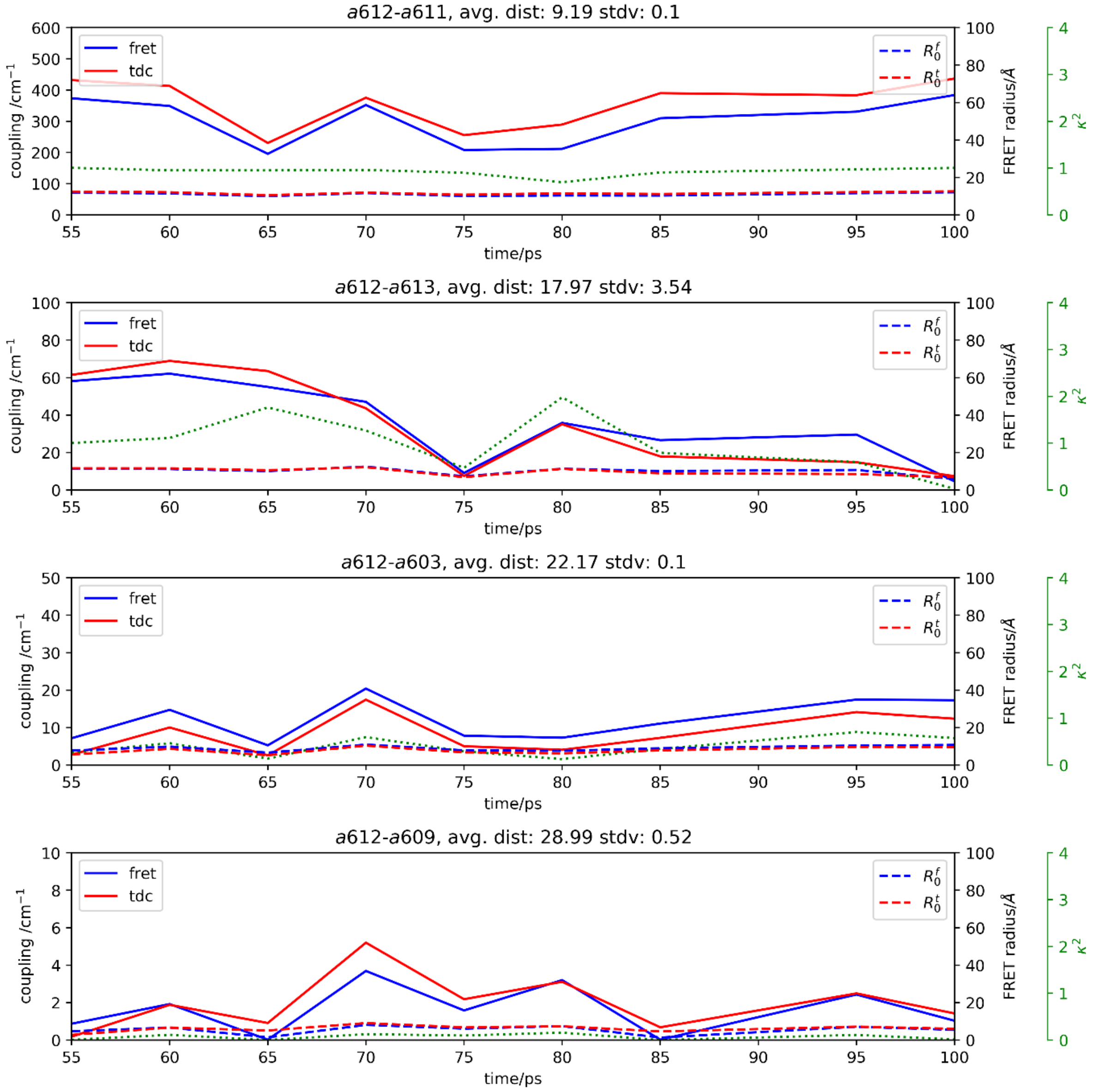


*Figure 2: $B_x$ couplings from the FRET and TDC methods and the corresponding FRET radii and $\kappa^2$ values, between different Chl a-a pairs (A-D).*
